# Sequential HIV-1 fusion protein gates control the initiation of host cell entry

**DOI:** 10.64898/2026.08.06.743292

**Authors:** Hung Do, Muralikrishna Lella, Devansh Fulmali, Niva Rajpara, Ilona Unarta, Alexis Johnson, Katarzyna Janowska, Helena Laukaitis, Carrie Saunders, R.J. Edwards, Priyamvada Acharya, Amit Sharma, S. Gnanakaran, Rory Henderson

## Abstract

HIV-1 fusion with host cells is initiated upon engagement of the viral envelope (Env) protein by its receptors and co-receptors. Receptor-induced conformational changes in the Env ectodomain expose the fusion machinery and position the HIV-1 fusion peptide for release and transit to the host membrane. However, no allosteric rearrangements in the envelope receptor/co-receptor binding domain that could induce fusion peptide release have been identified, and the trigger for the structural cascade that brings the viral and host membranes together remains unknown. Here, we identified two sequential conformational gates that control fusion peptide release, thereby facilitating its transition to the host membrane. We show that destabilizing a clasp holding the fusion peptide proximal motif enhances fusion peptide release, and that a coupled downstream gate controls the release of the receptor-binding gp120 subunit. Single-particle cryo-EM structures, biased and unbiased atomistic simulations, and functional experiments reveal that this process is controlled by shifts in the angular orientation of the gp120 subunits, induced by the geometries required for receptor and co-receptor binding. These results show how HIV-1 senses contact with the host cell to initiate fusion, thereby permitting viral entry.

## Introduction

The HIV-1 Envelope (Env) protein mediates host cell fusion through a series of receptor-induced conformational changes.^1^ Starting from a pre-fusion, pre-receptor, closed conformation that protects conserved components, Env transitions through a poorly understood structural intermediate, the pre-hairpin, that anchors the host and virus membranes.^2–4^ The pre-hairpin initiates virus-cell membrane fusion, leading to a stable post-fusion state composed of a 6-helix bundle.^5–7^ While considerable structural information exists describing pre-and post-fusion states, a structure-based understanding of pre-hairpin formation is limited.

The HIV-1 Env is a trimer of glycosylated heterodimers composed of a receptor-binding ∼120-kDa gp120 subunit and a 41-kDa gp41 subunit that contains fusion elements and anchors the protein to the viral membrane.^1^ The CD4 receptor on the surface of T cells triggers a series of conformational changes in gp120 that result in its rotation about the central Env trimer axis, exposing conserved portions of the gp41 fusion machinery.^8–18^ This includes extension of the central heptad repeat 1 (HR1) three-helix bundle and rearrangements in the disposition of the Env fusion peptide (FP) and the fusion peptide proximal region (FPPR).^4,8,10,18^ These changes in the trimer conformation are a prelude to the formation of the putative pre-hairpin intermediate that is characterized by the extension of the HR1 three-helix bundle.^4^

The FP is believed to be the first direct contact between the HIV-1 Env and the host membrane.^4^ The FP is exposed in the pre-fusion closed state and is anchored to the Env via FPPR interactions with gp120 and gp41 near the transmembrane trimer base.^8,10,19^ The FPPR leads through HR1 into the central three-helix bundle.^20^ Receptor-induced gp120 rotation results in FP sequestering in a gp120/gp41 groove and FPPR/HR1 helical extension toward the adjacent protomer.^8,10,14,21^ The FPPR nevertheless remains anchored to the trimer base.^8,10,14^ Pre-hairpin formation requires release of the FP/FPPR to enable three-helix bundle extension and FP transit to the host membrane, but this release has not been observed in high-resolution prefusion closed or open states.

The FPPR is embedded in a cleft formed between gp120 and gp41 in the prefusion closed state.^22^ The FPPR is positioned at the C-terminus of the FP, forming a gp120-interactive random coil-turn-helix motif that connects to a disordered HR1 segment (Figure 1A). Upon CD4 receptor activation, gp120 rotation induces FPPR rotation about the turn motif methionine, M530, extending the FPPR helix toward the adjacent protomer and permitting close FP interaction with the gp120/gp41 cleft (Figure 1A).^8^ This methionine acts as a central anchor for the FP/FPPR. It is held by a tryptophan clasp (W-clasp)^22,23^, which is an arrangement of three flanking gp41 tryptophans held in place by a helix-turn-helix motif in alpha helix 8 (alpha-8) (Figure 1A).^23^ The interaction of the FP with the gp120/gp41 cleft, the FPPR/HR1 contact with the adjacent protomer, and the M530-W-clasp engagement together inhibit pre-hairpin formation.

**Figure 1.**
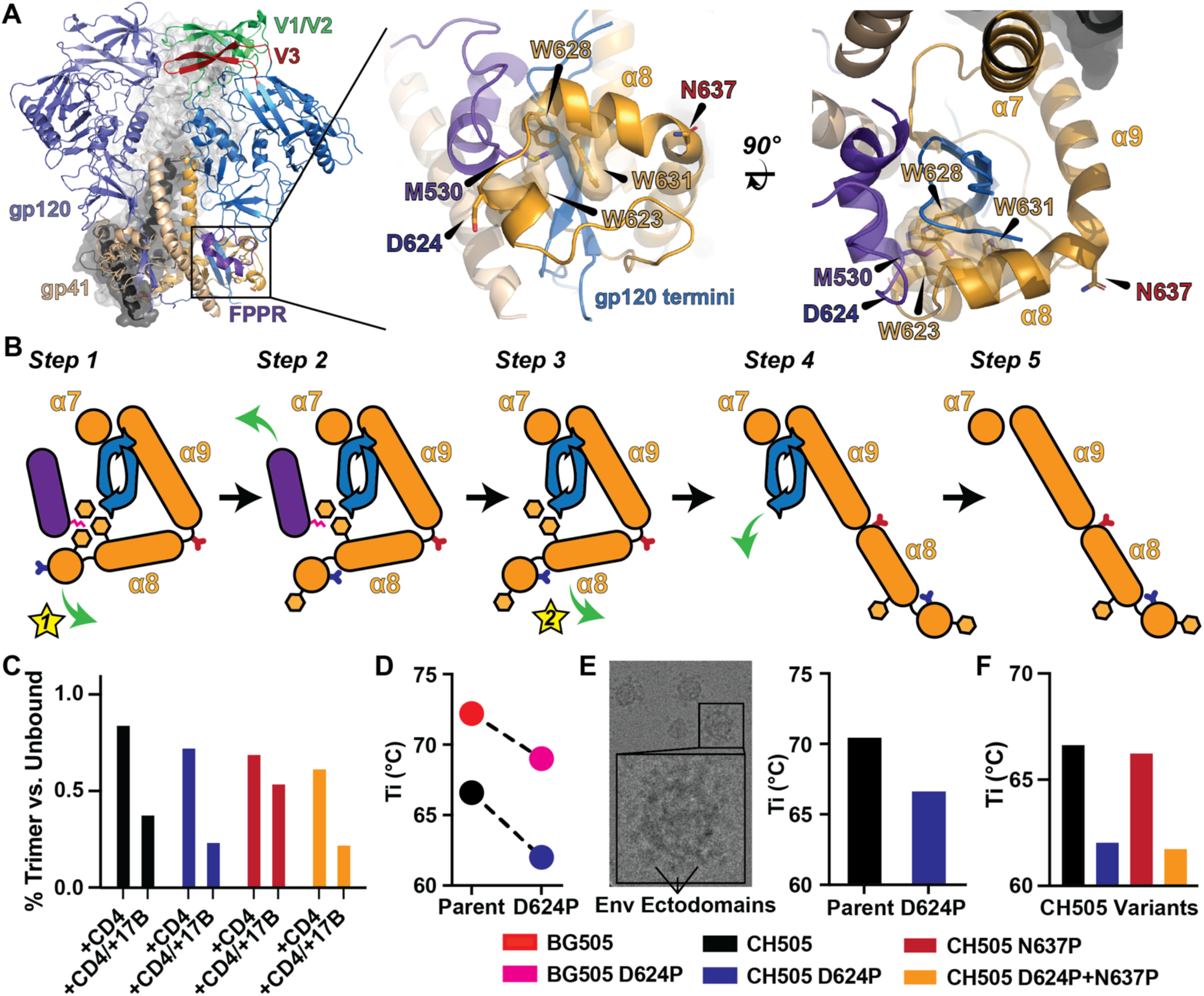
Two sequential structural gates control FPPR release and subsequent gp120 shedding. **A)** (left) A prefusion closed Env structure highlighting the receptor/co-receptor binding gp120 and fusion mediating gp41 domains. The trimer apex is formed by the V1/V2 and V3 hypervariable motifs (green and red, respectively). The trimer base consists of a gp41 collar that surrounds the gp120 termini, with the FPPR (purple) serving as a mobile segment that anchors the FP near the viral membrane. (right) The gp41 collar depicting FPPR M530 capture by the alpha-8 tryptophan-clasp. Gate perturbation sites are identified-at the W623 proximal D624 and the alpha-8-to-alpha-9 HR2 helix-turn-helix N637 positions. **B)** A cartoon representation of the proposed FPPR W623 gate 1 and gp120 termini HR2 helical extension gate 2. **C)** Receptor-and co-receptor-induced changes in trimer morphology as determined by NSEM 2D class average populations. Quantities are expressed as a fraction of total trimer particles identified in micrographs for ligand-free trimers vs. receptor CD4 and receptor CD4 plus co-receptor mimicking antibody 17b bound trimers. **D)** Thermal denaturation inflection temperatures (T_i_) determined using differential scanning fluorimetry (DSF) for the parent BG505 and CH505 constructs compared to the D624P constructs. **E)** (left) A representative cryo-EM micrograph depicting CH505 ESCRT, prefusion-stabilized gp150 trimer VLPs, highlighting Env ectodomains. (right) Inflection temperatures, T_i_s, determined by DSF for the CH505 parent and D624P VLP Envs. **F)** Inflection temperatures, T_i_s, determined by DSF comparing the CH505 parent ectodomain trimer to D624P, N637P, and combined D624P/N637P denaturation temperatures. Thermal denaturation inflection temperatures display production lot standard errors of 0.06-0.1 (Supplemental Figure 3E; N=2-12). One lot represents an independent construct transfection and purification.

A late pre-hairpin intermediate formed between the virus and CD4+ T cells bound to HR2-mimicking inhibitor peptides was visualized by cryo-ET.^2^ This intermediate showed long, tube-like structures connecting the viral and host cell membranes, but lacked clear densities for the gp120 domains or receptors.^2^ More recent cryo-ET studies of viral Env interacting with virus-like particles displaying the CD4 receptor showed that the Env and virus are drawn closer to the host membrane as multiple CD4 molecules engage the trimer.^24^ The Env displays an asymmetric open state that closely matches high-resolution CD4-bound, prefusion open-state trimer ectodomains.^12,24^ However, neither structure displays the degree of helical extension in HR1 expected for the pre-hairpin intermediate, nor is the FP/FPPR released from the trimer base. The crystal structure of gp120 bound to CD4 and CCR5 did not reveal an allosteric rearrangement related to co-receptor binding^25^. It is therefore unclear how receptor and co-receptor anchoring triggers FP/FPPR release to initiate host-cell fusion.

The FPPR M530 interaction with the alpha-8 tryptophan clasp is the sole invariant structural constraint that restricts FP/FPPR movement. Structural analysis of the clasp indicates that the distal W623 residue is uniquely positioned to move away from M530. Thus, we hypothesized that distal alpha-8 W623 contact with M530 gates FP/FPPR release. Release of the FP/FPPR eliminates a substantial portion of the gp41 contact with gp120 that anchors the receptor binding domain to the trimer. Molecular simulation further suggests that W623-gated release of the FP/FPPR results in subsequent clasp release of gp120 constraints, thereby facilitating gp120 shedding. We show that FP/FPPR release is governed by two sequential gates: destabilizing the W623 gate and inhibiting gp120-gated shedding. High-resolution structures and molecular dynamics simulations show that these gates are triggered by co-receptor binding through restrictions in trimer geometry that force the W623 gate open. Together, these results show how HIV-1 initiates fusion by sensing co-receptor binding.

## Results

### Two sequential structural gates control FPPR and gp120-termini exposure

The Env gp41 base is composed of FP/FPPR residues and post-HR1 residues 597-664.^22^ Residues 610-628 make up alpha-8 and contain the three FPPR M530 anchoring tryptophan residues at positions 623, 628, and 631 (Figure 1A).^22^ These motifs, together with HR2 residues, surround the gp120 termini, anchoring gp120 to the trimer. Release of the FPPR M530 requires tryptophan-clasp disengagement. The W628 and W631 residues pack against M530 and the gp120 termini, while W623 resides at the distal end of a helix, packing only against M530 (Figure 1A). The W623 alpha-8 turn motif therefore serves as a gate through which W623 releases M530 from clasp contact, thereby permitting FPPR release from the gp41 base. FP/FPPR release would, in turn, eliminate gp41 base closure around the gp120 termini, leaving the termini exposed to release and gp120 shedding (Figure 1B). We simulated 25 independent replicas of the gp120 termini and surrounding gp41 collar with the FP/FPPR absent. The atomistic simulations revealed that the remaining W628/W631 contact with the gp120 termini is rapidly broken (Supplemental Figure 1). This is followed by extension of the HR2 helix at an alpha-8 to alpha-9 helix-turn-helix motif at position 637 (Supplemental Figure 1). These structural transitions represent a two-gate system that controls release of the FP/FPPR and subsequent gp120 shedding.

### Destabilization of the W623 gate induces premature Env trimer disassembly

Prefusion-stabilized trimers cannot shed gp120, access the pre-hairpin, or transition to post-fusion structural states^26,27^, yet remain subject to receptor-and antibody-mediated disassembly^28^. We hypothesized that FPPR release is a gateway to trimer disassembly owing to its essential role in completing the gp120 termini collar. Since the FPPR is anchored to the collar via M530 capture by the tryptophan clasp, we reasoned that disrupting the clasp would render the trimer sensitive to FPPR release, thereby inducing disassembly. To test this hypothesis, we introduced a proline mutation at position D624 in the distal tryptophan-clasp W623 gate in two prefusion-stabilized trimer ectodomains. These included a clade A BG505 isolate^27^ and a chimeric trimer pairing the BG505 gp41 with a clade C CH505^29^ transmitted founder gp120.^30^ The CH505 transmitted founder chimera (CH505) contained additional mutations that disable CD4-induced allosteric rearrangements in gp120^31^ (F14) and two additional prolines^32^ in HR1 that enhance expression. The purified constructs showed little change in conformation-dependent antibody binding, indicating that the populations of closed and open states were minimally perturbed (Supplemental Figure 3A-C). The prefusion-stabilized trimer ectodomains both contain a disulfide between the gp120 N-terminus and gp41 base.^27^ This prevents complete separation between gp120 and gp41 but does not prevent FP/FPPR release. We first asked whether engagement of the receptor and co-receptor mimic 17b antibody disrupts trimer structural morphology in the D624P constructs. Negative-stain electron microscopy revealed that CD4 and CD4/17b binding sequentially induce ectodomain disassembly, thereby reducing the overall populations of well-folded trimers (Figure 1C, Supplemental Figure 2). This effect was more pronounced in the CH505 D624P construct (Figure 1C, Supplemental Figure 2). These results indicate that the D624P mutation can enhance receptor-induced disassembly.

The prefusion-stabilized BG505 ectodomain trimer exhibits a single thermal denaturation transition, in contrast to the biphasic melt of monomeric gp120, indicating that the trimer denatures as a single unit.^27^ Given Env trimer denaturation is irreversible, these observations are consistent with denaturation controlled by a kinetic gate.^33^ If FPPR release acts as a kinetic gate for Env ectodomain denaturation, premature FPPR release mediated by destabilization of the D624P W623 gate would result in a reduced thermal denaturation temperature. We measured thermal denaturation temperatures for each trimer using differential scanning fluorimetry (DSF). Both trimer construct parents and D624P mutants displayed a single, symmetric denaturation peak (Supplemental Figure 3). The D624P BG505 trimer displayed a 3.2 °C thermal inflection (T_i_) reduction relative to the parent 72.2 °C T_i_ (Figure 1D, Supplemental Figure 3). Similarly, the D624P CH505 chimera displayed a 4.6 °C reduction relative to the parent 66.6 °C T_i_ (Figure 1D, Supplemental Figure). These effects extended to virus-like particle membrane-embedded Envs containing the transmembrane motif (Figure 1E). Enhanced receptor/co-receptor mimic and temperature-induced disassembly in the D624P constructs are consistent with premature FPPR release.

### An HR2-gate limits receptor/co-receptor antibody mimic-induced Env trimer disassembly

The gp41 collar structure allows W-clasp W623-mediated release of the FPPR M530 residue. Release of the FPPR would expose a substantial portion of the gp120 termini to solvent, eliminating key contacts that anchor the gp120 subunit to the trimer. The remaining gp41 W-clasp W628 and W631 residue contact with the termini would nevertheless block gp120 release. The simulations showed that extension of the HR2 helix eliminated these W-clasp contacts with the gp120 termini (Supplemental Figure 1). The HR2-gate is composed of an alpha-8-turn-alpha-9 helical motif. We hypothesize that introducing a proline residue at position N637 in the turn would stabilize the turn motif, prevent HR2 helical extension, and thereby inhibit trimer disassembly. The purified CH505 N637P mutation had minimal impact on conformation-dependent antibody binding, showing a roughly twofold reduction in V3-loop-targeting antibody 19b^34^ (Supplemental Figure 3A and B). This suggests the mutation has a minor stabilizing effect on the prefusion-closed conformation. We next asked whether the N637P mutation impacts receptor-and co-receptor-induced trimer disassembly. The N637P mutation did not protect the trimer from CD4 receptor-induced disassembly but did protect it from disassembly induced by binding the co-receptor mimic 17b antibody (Figure 1C, Supplemental Figure 2). This is consistent with a disassembly protective effect for alpha-8-turn-alpha-9 stabilization. Unlike the W623 gate D624P substitution that directly impacts the gate holding the FPPR in place, the N637P substitution is not predicted to affect FPPR release. Consistent with this prediction, the N637P mutation did not affect the trimer thermal denaturation inflection temperature (Figure 1F, Supplemental Figure 3H and I). We next asked whether the N637P substitution impacts the D624P substitution-induced effects on trimer disassembly. The purified CH505 D624P+N637P construct displayed conformation-dependent antibody binding similar to the parent CH505 construct, abrogating the minor CH505 N637P-induced close-state stabilization (Supplemental Figure 3A and B). The disassembly assay revealed a reversion to the CH505 D624P CD4 and CD4/17b-induced disassembly phenotype, eliminating the protective effect observed for the CH505 N637P CD4/17b-induced disassembly (Figure 1C, Supplemental Figure 2). Similarly, the thermal denaturation profile for the CH505 D624P+N637P construct matched the CH505 D624P construct (Figure 1F, Supplemental Figure 3H and I). These results show that the N637P mutation can protect against trimer disassembly but cannot restore the ectodomain FPPR release phenotype induced by the D624P substitution.

### W623 gate and HR2 gate destabilization impair virus fusion

While the prefusion-stabilized Env trimer ectodomains permit interrogation of transitions between the prefusion closed and open states^8,10,12,17^, the gp120-to-gp41 disulfide linkage prevents the examination of downstream events, including gp120 shedding^26^. We therefore asked whether gate disruption affects Env-mediated fusion of native viruses with host cells. We hypothesized that W623 and HR2 gates play essential, coupled roles in regulating the fusion process. The W623 gate senses co-receptor engagement, initiating the fusion event. The HR2 gate then regulates gp120 release, ensuring the virus remains anchored to the host through receptor interaction while the FP transits to the host membrane. Premature FPPR release and inhibition of gp120 shedding induced by gate disruption would interfere with these fusion processes, thereby impacting viral infectivity. We therefore first assessed the effects of the D624P and N637P mutations on HIV-1 infectivity. Introduction of the D624P mutation resulted in a significant reduction in viral titers relative to wild-type (WT) virus (Figure 2A). Similarly, the N637P mutation also led to a marked decrease in infectivity. Although the magnitude of this defect was comparable, it was mechanistically distinct (Figure 2A). When combined, the D624P+N637P double mutant exhibited further reductions in infectivity relative to WT (Figure 2A, Supplemental Figure 4). We next evaluated whether these infectivity defects correlated with impaired membrane fusion. The HIV-1 Blam-Vpr fusion assay showed that both D624P and N637P mutants induce substantial reductions in fusogenicity compared to WT Env (Figure 2B, Supplemental Figure 4). These findings indicate that both mutations impair Env-mediated virus-cell membrane fusion, consistent with their reduced infectivity phenotypes. These results show that prolines in the W623 gate and the HR2 gate significantly impair virus-host cell fusion.

**Figure 2.**
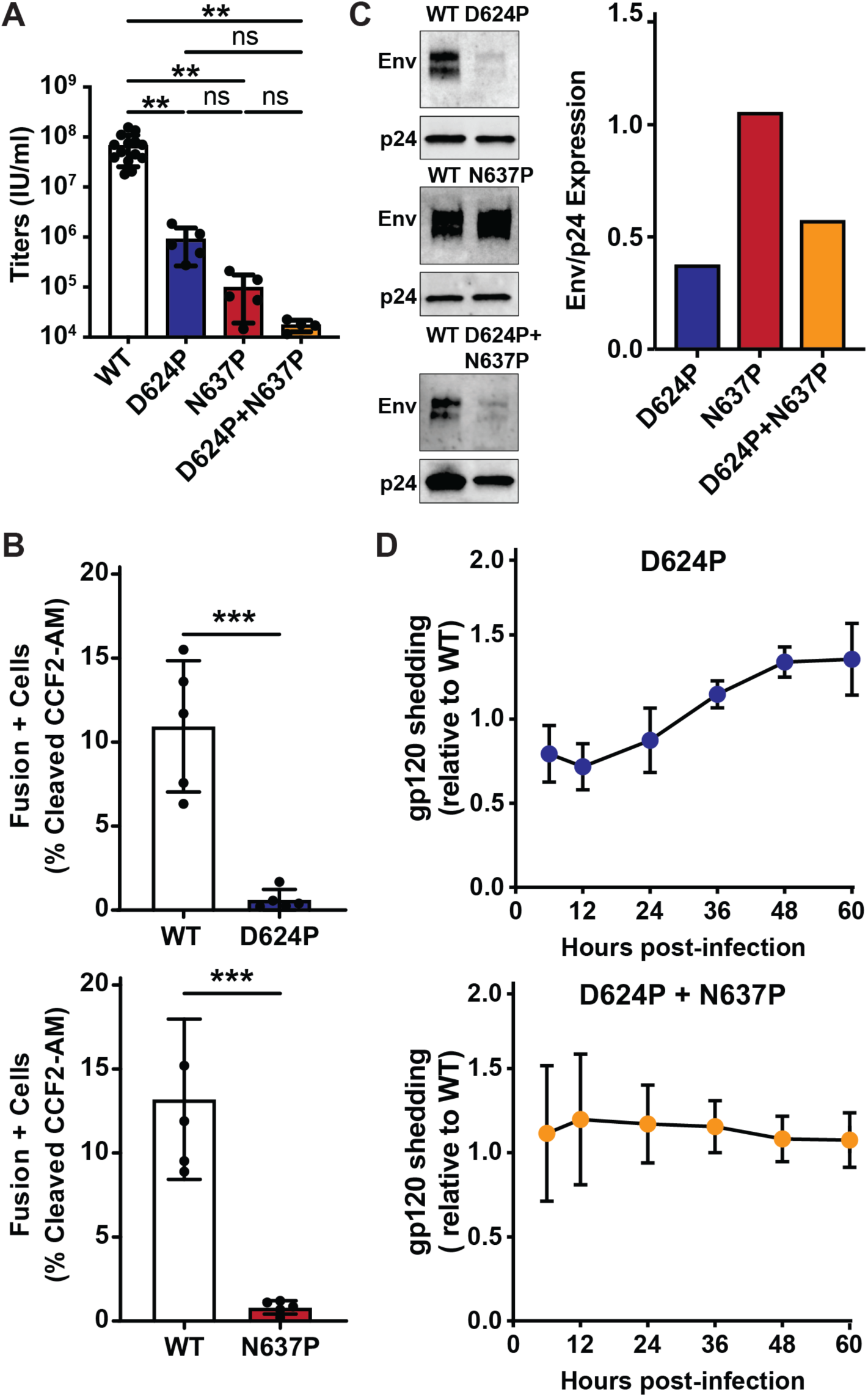
Blocking HR2 helical extension rescues the wild-type shedding phenotype from induced W623 gate instability. **A)** TZM-bl infectivity assay of HIV-1 produced from HEK293T cells transfected with WT or mutant Env proviral constructs. Viral titers are shown as infectious units per milliliter (IU/ml) on a logarithmic y-axis, with the corresponding WT or mutant Env proviruses on the x-axis. Bars represent the mean of n = 14, 5, 5, and 4 independent experiments for WT, D624P, N637P, and D624P+N637P, respectively; individual points represent single experiments and error bars indicate standard deviation. All pairwise comparisons were performed by ordinary one-way ANOVA with Tukey’s multiple comparisons test (**p = 0.0020 for WT vs. D624P; **p = 0.0017 WT vs. N637P; **p = 0.0040 for WT vs. D624P + N637P; p > 0.9999 for all comparisons among mutants, ns). **B)** Quantification of viral fusion for the indicated HIV-1 variants (x-axis), presented as percent fusion. Data represent the mean of five independent experiments, with error bars indicating standard deviation. Statistical comparisons between WT and mutant Env variants were performed using an unpaired, two-tailed *t* test (***p = 0.0004 for WT vs. D624P and WT vs. N637P). **C)** Western blot analysis of virions (*left*) harvested at 48 hours post-transfection from HEK293T cells transfected with the indicated HIV-1 proviral plasmids. Immunoblotting performed using anti-Env and anti-p24 antibodies. Quantification of the Env-to-p24 ratio (*right*). **D)** Quantification of gp120 shedding over time. The y-axis represents gp120 levels in post-ultracentrifugation supernatants from indicated mutant HIV-1 normalized to wild-type (WT), the x-axis indicates the time points post-transfection at which supernatants were collected. gp120 levels were determined by densitometric analysis of immunoblots. Data represent the mean of five independent experiments, with error bars indicating standard deviation.

### The W623 and HR2 gates control sequential FPPR release and gp120 shedding

Impaired virus fusion by the W623 and HR2 proline substitutions is expected based on their putative gating roles in FP/FPPR release and gp120 shedding. The prefusion-stabilized ectodomain disassembly assays show that the W623 gate D624P substitution enhances disassembly, while the HR2 gate N637P substitution limits disassembly. The combined substitutions, however, revealed that N637P did not protect from the enhanced D624P-induced disassembly phenotype. The pre-fusion stabilization mutations introduce a disulfide between gp120 and gp41 to prevent gp120 shedding. Thus, these assays to not fully capture the shedding phenotype. We hypothesized that, absent covalent linkage between gp120 and gp41 in virions, the D624P substitution would show an enhanced shedding phenotype. Conversely, we hypothesized the N637P substitution would not display a shedding phenotype, but could recover D624P-induced shedding by limiting gp120 termini exposure. We first quantified Env levels in both producer cells and purified virions. The D624P mutation resulted in a pronounced reduction in virion-associated Env, despite relatively modest effects on cellular Env expression (Figure 2C). This suggests that D624P disrupts Env stability or incorporation into virions. In contrast, the N637P mutation did not significantly alter either cellular or virion Env levels (Figure 2C), indicating that its defect in infectivity and fusion arises independently of Env incorporation. The D624P+N637P double mutant partially restored virion Env levels compared to D624P alone (Figure 2C, Supplemental Figure 4), suggesting a compensatory interaction between these residues. Despite this partial rescue of Env incorporation, infectivity remained impaired, indicating that restoration of Env levels alone is insufficient to fully recover function. Given the reduced virion Env levels observed for D624P, we next examined whether this phenotype could be explained by increased gp120 shedding.^35,36^ Time-course analysis revealed that the D624P mutant exhibited elevated gp120 shedding relative to WT (Figure 2D), consistent with Env trimer destabilization and loss of gp120 from the virion surface. The D624P+N637P double mutant displayed reduced gp120 shedding compared to D624P alone, with the effect becoming more pronounced at later time points (Figure 2D). These results indicate that while D624P destabilizes Env and promotes gp120 shedding, N637P can partially counteract this effect in the double mutant, consistent with the W623 and HR2 gated mechanism controlling FP/FPPR release and gp120 shedding, respectively.

### Env trimers display asymmetric W623 gate structural heterogeneity

The ectodomain disassembly and virion Env shedding patterns indicate that W623-gated FPPR release and subsequent HR2-mediated helical extension lead to gp120 shedding while presumably freeing the FP for transit to the host membrane. It is not clear, however, what induces the W623 gate to release FPPR M530 upon receptor engagement at the host membrane. Recently published cryo-ET data showed that sequential binding of CD4 molecules brings the virus closer to the host membrane.^24^ The co-receptor binding site nevertheless remains too far from the membrane to engage CCR5.^24,25^ Further, a gp120-bound CCR5 structure did not identify allosteric rearrangements leading from the co-receptor binding site to the gp120 termini.^25^ The gp41 collar is known to shift positions in response to receptor-binding-induced changes in gp120 tilt.^31^ We therefore asked whether the W623 gate structure is impacted by gp41 position. We examined the tryptophan-clasp conformation in published high-resolution prefusion closed and open structures of the BG505 Env ectodomain.^12,37^ The prefusion-stabilized Env ectodomain structure of the BG505 HIV-1 isolate was determined in the presence and absence of CD4^37^. These include ligand-free and single-CD4-bound prefusion closed structures, as well as an asymmetric 2-CD4-bound open state^12^ (Figure 3A, Supplemental Figure X). We first examined the structures of the ligand-free and single-CD4-bound prefusion closed states, as well as the two CD4-bound open states. The secondary and tertiary structures of the clasps within and between each structure were similar (Supplemental Figure 5). However, clear differences in tryptophan-clasp resolvability between protomers within each structure were visually apparent (Figure 3B, Supplemental Figure 5). Differences in resolvability often result from conformational heterogeneity.^38^ In the case of pre-fusion closed states, ligand-free BG505 and single-CD4-bound BG505, the overall conformations of the FPPR and α8 regions were retained; their resolvability differed among protomers and was more evident in the 3^rd^ protomer. In the two-CD4-bound open state, only one protomer clasp retained clear secondary-structure resolvability, and quantitative analysis confirmed that this asymmetric loss of density is consistent with local conformational heterogeneity during CD4-induced Env opening (Figure 3B, Supplemental Figure 5). These resolvability features were recapitulated using quantitative metrics and were independent of overall protomer resolvability (Supplemental Figure 5). Thus, while the overall tryptophan clasp conformation is retained, it exhibits asymmetric protomer positional heterogeneity that depends on trimer conformation.

**Figure 3.**
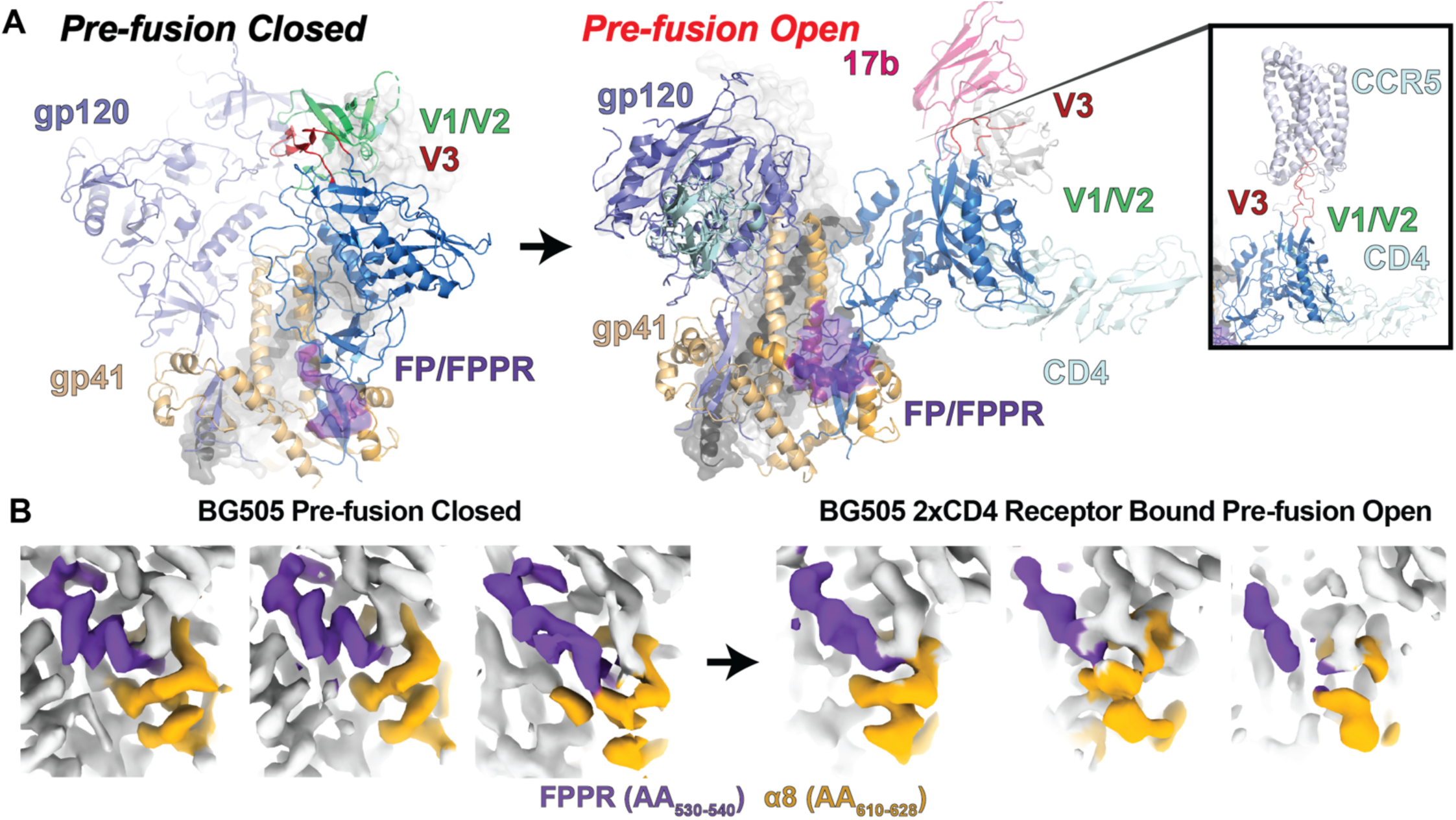
The W623 gate is structurally heterogeneous. **A)** (left) The prefusion closed Env trimer ectodomain structure highlighting the FP/FPPR conformation at the trimer base. (right) The CD4-and co-receptor-mimicking 17b antibody bound to the Env open structure, highlighting the shift in FP/FPPR conformation. (inset) The co-receptor CCR5 bound to gp120, depicting the interaction between the gp120 V3 loop and exposed V1/V2 motif. **B)** Maps reconstructed from single-particle cryo-EM showing the FPPR (purple) and alpha-8 (yellow) densities for each protomer in the ligand-free BG505 prefusion closed state (left) and the two-CD4-bound prefusion open state (right).

### The gp41 tilt angle is a function of the gp120 tilt angle

The observed tryptophan-clasp structural heterogeneity shows the clasp is responsive to differences in Env conformation. Co-receptor binding requires a closer Env approach to the host membrane that could impact the degree of gp120 rotation relative to the gp41 core. If gp120 rotation impacts the tryptophan clasp structure, co-receptor-induced rotation could act as an allosteric switch that triggers FP/FPPR release. We therefore asked whether gp120 rotation impacts the tryptophan clasp geometry. We define gp120 rotation as a tilt angle using a previously identified hinge point near the gp120 termini at the trimer base (Figure 4A).^18^ The gp120 rotation was accompanied by rotation of the gp41 collar. We defined gp41 rotation as the tilt of the clasp alpha-8 helix motif within the plane of the trimer base (Figure 4A). For this analysis, we included the ligand-free and CD4-bound BG505 structures as well as a previously published ligand-free 1059 isolate^37^ prefusion closed-state trimer. Additionally, we obtained single-particle cryo-EM reconstructions of the ligand-free CH505 prefusion-stabilized trimer, resolving prefusion closed and open structures at global resolutions of 3.97 Å and 6.35 Å, respectively (Supplement Figures 6 and 7). The gp120 and gp41 tilt angles were determined from cryo-EM map-fitted structures for each protomer in each construct. The angles revealed substantial protomer-tilt asymmetry in both the prefusion closed-and open-state structures. A linear regression fit of gp120 tilt vs. gp41 tilt showed a significant linear relationship between the two angles, with a Spearman’s rho of 0.66 (Figure 4A; R^2^=0.44, p=0.004). These results show that gp120 and gp41 rotation are correlated, providing an allosteric path between distal gp120 contact and the FPPR anchor at the base of gp41.

**Figure 4.**
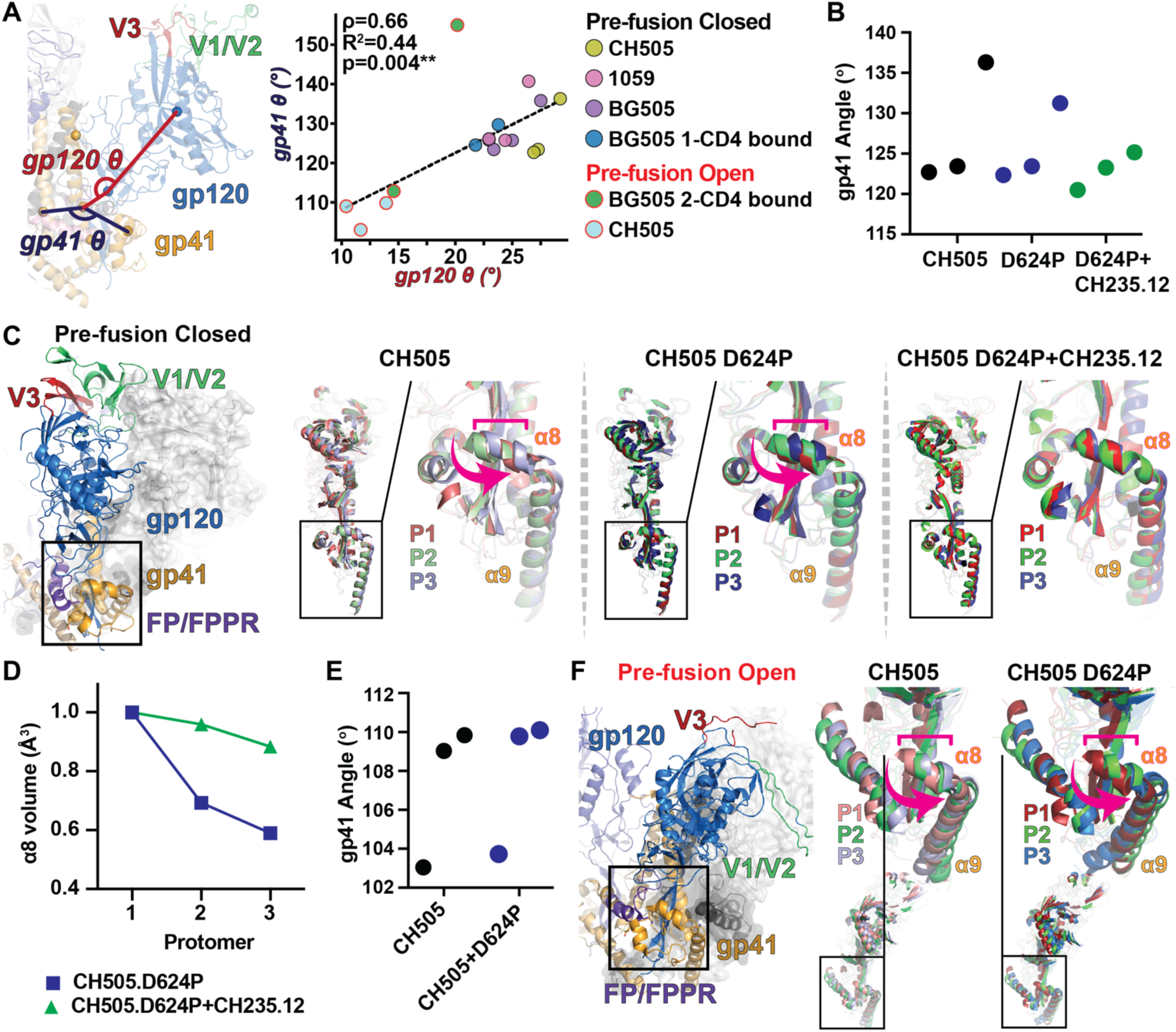
Receptor-binding domain gp120 tilt correlates with gp41 angle and W623 gate structural heterogeneity. **A)** (left) Representative open-state gp120 protomer highlighting the gp120 and gp41 tilt angle measurements. (right) gp120 vs. gp41 tilt angles of protomers in ligand-free and CD4-bound receptor structures. The dashed line indicates the linear regression fit to the data. **B)** The prefusion closed-state protomer gp41 tilt angles for the CH505 parent, D624P, and CD4 receptor-binding-site-targeting CH235.12 broadly neutralizing antibody-bound D624P constructs. **C)** (left) A prefusion closed-state trimer structure highlighting the alpha-8/9 and FPPR region. (right) Zoomed-in views of the helix/sheet motifs in the alpha-8/9 and FPPR regions of the CH505 parent, D624P, and D624P+CH235.12 prefusion closed-state structures. The pink arrows and brackets indicate the direction of rotation of gp41 between the protomer alpha-8 helices. **D)** The normalized resolvable alpha-8 volumes for each protomer in the CH505 parent, D624P, and D624P+CH235.12 maps. **E)** (left) A prefusion open-state trimer structure highlighting the alpha-8/9 and FPPR region. (right) Zoomed-in views of the helix/sheet motifs in the alpha-8/9 and FPPR regions of the CH505 parent and D624P prefusion open-state structures. **F)** The prefusion open-state protomer gp41 tilt angles for the CH505 parent and D624P structures.

### W623 gate structural heterogeneity tracks gp120 and gp41 tilt

The correlation between the gp120 and gp41 tilt angles indicates that clasp geometry varies with overall trimer geometry. The observed positional heterogeneity in the BG505 cryo-EM maps indicates that these geometric changes affect the clasp’s stability in holding the FPPR. The ligand-free, prefusion closed state structures each displayed a single protomer with a distinct gp41 angle relative to the other two. Examination of the map densities for the BG505 and CH505 revealed that the protomer with the most extreme gp41 tilt angle corresponded to the most conformationally heterogeneous alpha-8 clasp (Supplemental Figures 8 and 9). All protomers in the 1059 map displayed resolvable secondary structures in alpha-8, despite similar gp41 tilt asymmetry (Supplemental Figures 8 and 9). This trimer exhibits markedly greater gp120 scissoring motion^39^ compared to BG505.^37^ Examination of the gp120 angles indicated that the asymmetric protomer’s gp120 tilt favors interaction with the FPPR, thereby stabilizing the gp41 base in this protomer (Supplemental Figures 8 and 9). This observation is consistent with the correlation between gp120 and gp41 tilt angles, which shows that gp41 tilt is not a strict function of gp120 tilt angle. Thus, the clasp structural heterogeneity is a function of both gp41 tilt and FPPR structural stability.

### The W623-gate D624P mutation destabilizes the distal clasp helix

We hypothesized that W623 release of the FPPR depends on gp41 tilt, but that alpha-8 helix disruption inhibits the conformationally protective role tilt plays in clasp stability. We obtained single-particle cryo-EM reconstructions of the ligand-free CH505 D624P prefusion-stabilized trimer, resolving prefusion closed and open structures at resolutions of 3.52 Å and 7.48 Å, respectively (Supplement Figures 10 and 11). To test whether alpha-8 destabilization fully decouples the clasp from gp41 tilt, we also obtained a CH505 D624P prefusion closed-state reconstruction for a CH235.12^29^ broadly neutralizing antibody (bnAb)-bound structure. The CH235.12 bnAb binds to the CD4 receptor binding site. Antibodies targeting this site are known to stabilize the closed state of viral Env^40^ and often yield symmetric prefusion-stabilized Env ectodomain reconstructions.^41^ The CH235.12-bound D624P reconstruction allows us to determine whether induced gp120 and gp41 tilt symmetry leads to symmetry in the structural heterogeneity of the clasp, despite W623-gate instability. We obtained a single-particle cryo-EM reconstruction of the CH235.12-bound CH505 D624P trimer, yielding a single prefusion closed map at 4.16 Å (Supplement Figure 12). The gp120 and gp41 tilt angles of the D624P structure showed similar asymmetry to the parent CH505 structure (Figure 4B). As expected, the CH235.12-bound D624P structure showed substantially reduced tilt asymmetry (Figure 4B). Examination of the ligand-free structures showed that the asymmetric protomer alpha-8 helix is shifted away from the FPPR (Figure 4C). The CH235.12-induced symmetry positions each alpha-8 helix toward the FPPR (Figure 4B and C). We asked whether these shifts in gp41 tilt correspond with changes in clasp structural heterogeneity. The ligand-free D624P shows a distinct heterogeneity pattern relative to the parent construct, with two protomers having distinct gp41 tilt angles, resulting in substantially greater heterogeneity than a single, more stable clasp (Figure 4D, Supplemental Figure 13). This shows that the disruption of the distal W623 gate inhibits gp41 tilt control. This inhibition is not complete, as the symmetry induced by the CH235.12 antibody results in a dramatic shift toward greater homogeneity in the clasp structure (Figure 4D, Supplemental Figure 13). These results show that the D624P mutation induces local clasp instability that responds to gp41 tilt. Together with the other ligand-free and CD4-bound receptor structures, these results demonstrate that the clasp that holds the FPPR and fusion peptide is structurally linked to the gp120 geometry.

### The prefusion open state structures are shifted toward FPPR protective gp41 angles

The fusion peptide is solvent-exposed and typically poorly resolved in prefusion closed-state Env ectodomains but becomes sequestered in a cleft between gp120 and gp41 in the prefusion open state.^8,10,12^ This rearrangement is accompanied by a shift in the HR1 helical segment between residues 531-547 away from the gp120/gp41 cleft, toward contact with the adjacent protomer. The gp120 rotation away from the trimer’s central axis exposes the internal HR1 three-helix bundle, eliminating steric hindrance to pre-hairpin formation, but shifts the fusion peptide toward a protective, captured state. We asked whether FP sequestering is accompanied by reinforced FPPR capture by the tryptophan clasp. Lower gp41 tilt angles were associated with reduced alpha-8 and FPPR structural variability in prefusion closed state structures. The open Env cryo-EM reconstructions had global resolutions poorer than 4.5 Å, with insufficient local map quality around the atomic models; therefore, these structures were excluded from the quantitative map-based analysis. The map resolutions were, however, sufficient for structure modeling. We therefore asked whether the gp41 tilt angles shifted toward lower, FPPR-protective angles. Like the ligand-free prefusion close state structures, the ligand-free CH505 and CH505 D624P prefusion open state structures displayed marked asymmetry in the gp41 tilt angles (Figure 4E). These angles were shifted lower compared to the prefusion closed conformation. Unlike the prefusion closed structures, whose asymmetry was characterized by two protomers with lower gp41 tilt angles relative to a single protomer with higher tilt angles, the prefusion open state structures displayed a single protomer with a lower gp41 tilt angle relative to two protomers with higher tilt angles (Figure 4E). The lowest-angle protomer in each structure displayed an alpha-8 helix shifted toward the FPPR, directionally consistent with prefusion closed state structures. These results are consistent with gp41 structural rearrangements that protect from FPPR release in the absence of a protective gp120 tilt angle.

### The gp120 and FPPR are structurally dynamic

The Env open state retains the FP/FPPR through a combination of FP interaction with the gp120/gp41 cleft and alpha-8 stabilization of the FPPR.^8^ The structure analysis shows that FPPR structural heterogeneity correlates with rotation in gp120 and gp41. We hypothesized that this structural heterogeneity results from dynamics within the FPPR and gp120/gp41 rotations. We used an enhanced-sampling molecular dynamics simulation technique to monitor FP/FPPR dynamics as a function of gp120 and gp41 tilt angles in a ligand-free B41 isolate open intermediate-state Env trimer ectodomain^42,43^. This approach, termed Gaussian-accelerated molecular dynamics^44^ (GaMD), introduces a boosting potential to the entire system to accelerate dynamic conformational sampling. We monitored the distances between W571, positioned at the distal end of the gp120 three-helix bundle relative to the FPPR, and either gp120 or the FPPR. The W571-gp120 distance reports gp120 rotation, and the FPPR distance reports FPPR movement directed toward the putative pre-hairpin conformation. Across three independent boosted simulations, we observed gp120 and FPPR structural dynamics (Figure 5A, Supplemental Figures 14 and 15, Supplemental Table 3). This resulted in a single flexible conformation with no discernable correlation (Figure 5A and B, Supplemental Figures 14 and 15, Supplemental Table 3). We next examined the structure of the W623 gate, measuring the distance between the W623 and FPPR M530 residue alpha carbons. Consistent with our open-state structure observations, the W623-M530 dynamics were asymmetric, with one protomer occupying a stable distance consistent with the initial, closed-state distance (5.1 Å), and two protomers displaying greater distance variance, resulting in partial gate opening (Supplemental Figure 15, Supplemental Tables 3 and 4). These results show that structural heterogeneity in gp120, FPPR, and W623 gate arises from continuous dynamics, rather than discrete, sharp transitions.

**Figure 5.**
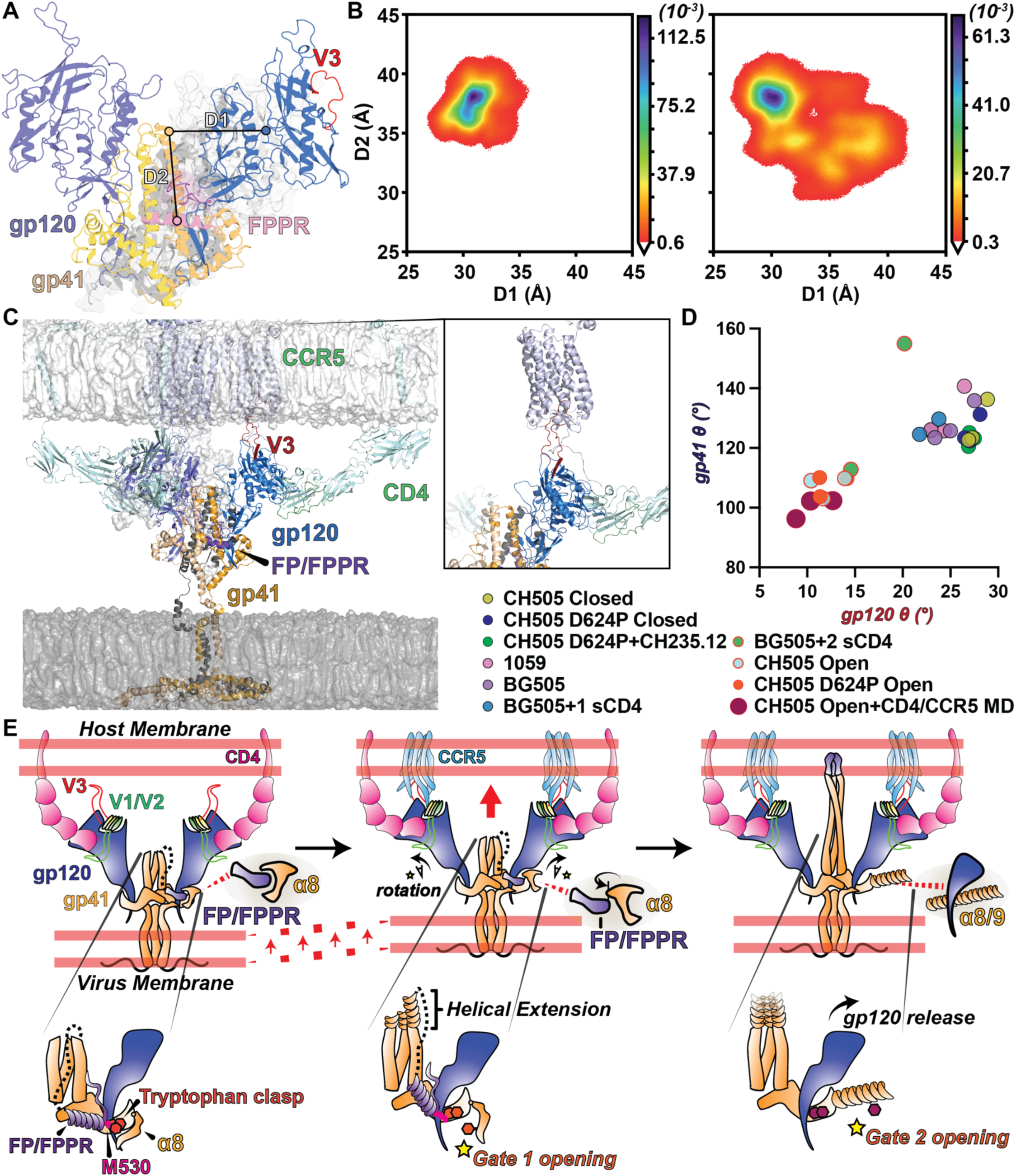
Receptor-binding domain gp120 tilt induces gp41 tilt, which induces FP/FPPR release. **A)** A representative wide-open state structure observed in the B41 open-intermediate LiGaMD enhanced sampling simulations identifying the W571-to-gp120 centroid (D1) and W571-to-FPPR centroid (D2) distance metrics. **B)** D1 vs. D2 kernel density estimates for the GaMD (left) and FP LiGaMD3 (right). **C)** The membrane-embedded CH505 Env gp160 bound to the membrane-embedded CD4 and CCR5. **D)** gp120 vs. gp41 tilt angles for the ligand-free and CD4-bound structures, including the CH505 D624P and simulated CD4/CCR5 bound gp160 membrane system. **E)** Cartoon representation of the proposed mechanism for FP/FPPR release and pre-hairpin intermediate formation.

### Enhanced FP dynamics induce greater gp120 rotation

The B41 open-intermediate GaMD simulations showed that the ligand-free Env ectodomain protomers display dynamic domain and secondary structural rearrangements consistent with the cryo-EM structure data. However, we did not observe FP or FPPR transit toward the trimer apex. The ligand-free 1059 structure showed that gp120 rotation toward the FPPR can stabilize the gp41 collar. The open state structures show that, when CD4 induces gp120 rotation away from the FPPR, the FP buries in a cleft between gp120 and gp41.^8,10,12^ This is accompanied by a shift in gp41 rotation. We hypothesized that these rearrangements act to protect the FP from premature release. We therefore asked whether adding a large boost potential to the FP residues specifically could initiate FPPR release. We introduced a large boosting potential to FP residues 512-522 in each B41 protomer using ligand GaMD (LiGaMD^45^) (Figure 5B and D, Supplemental Figures 16-17, Supplemental Tables 2 and 4). The LiGaMD3 simulation differed markedly from the GaMD simulations. The W571-to-gp120 distance showed marked asymmetry, with a single protomer, protomer 3 in each, shifting toward greater distances across all three independent simulations (Figure 5B, Supplemental Figures 16, 18 and 19, Supplemental Tables 4 and 5). This increased distance was associated with a reduction in the distance between the FPPR and W571 and, in one simulation, a discrete W623-gate opening event. These transitions involved an extensive network of residue contacts between gp41 and gp120 residues, consistent with large-scale steric perturbation of the gp41 collar fold (Supplemental Table 6). Together, these results show that destabilization of the FP contact with the gp120/gp41 cleft leads to increased gp120 rotation that initiates W623 gate opening and FPPR transit.

### Membrane-embedded co-receptor binding shifts gp120 and gp41 tilt angles

Recent cryo-ET of a virus interacting with membrane-embedded CD4 shows that, as additional CD4 molecules bind to the Env, the virus shifts closer to the CD4 membrane.^24^ The viral membrane is not, however, within the necessary distance to allow FP interaction with the apposed membrane. The Env V3 binding site on CCR5 is proximal to the host membrane. CCR5 binding therefore closes the gap.^25^ However, how FP release from the gp120/gp41 cleft is triggered is not known. The LiGaMD3 simulations showed that FP stability in the gp120/gp41 cleft is essential for maintaining the gp41 collar, and the structural and simulation data show that gp41 dynamics correlate with gp120 rotation. The gp120 rotation is therefore a potential allosteric trigger for FPPR release. We hypothesized that co-receptor CCR5 binding not only brings the trimer closer to the host membrane, but also shifts the gp120 rotation angle toward the release trigger angle observed in the LiGaMD3 simulations. To test this hypothesis, we modeled a membrane-embedded, full-length CH505 gp160 trimer bound to three CD4 and three CCR5 structures embedded in an apposed membrane (Figure 5C). This model structure was then relaxed using molecular dynamics simulation across four independent unbiased simulations. Like the GaMD and LiGaMD3 simulations, the gp120 and gp41 subunits displayed structural dynamics and, consistent with the open state structures, showed asymmetry in the gp41 tilt angles (Supplemental Figures 21 and 22, Supplemental Table 7). The angles were, however, lower than those observed in the available structures, with the W571-to-gp120 centroid distance shifted toward the LiGaMD FPPR-triggered distance. These results support a model of allosterically induced FP/FPPR release that is triggered by CCR5 binding, inducing gp120 rotation that in turn rotates gp41, initiating W623-gated release of the FPPR M530, and subsequent HR2-gated release of gp120 (Figure 5E).

## Discussion

Here, we identified two sequential structural gates in the HIV-1 Env gp41 base that control the initiation of host-cell fusion. Destabilization of the distal tryptophan-clasp gate at W623 by the D624P mutation induced premature FPPR release, supported by reduced thermal stability, enhanced ectodomain disassembly induced by receptor and co-receptor mimics, elevated gp120 shedding from virions, and loss of infectivity. Blocking HR2 helical extension with N637P reduced co-receptor mimicking antibody-induced disassembly but did not protect against W623-gate destabilization in the disulfide-linked gp120/gp41 context. It did, however, suppress shedding in the D624P background, placing HR2 extension downstream of FPPR release. High-resolution structures showed that the clasp is not a static element. It exhibits protomer-asymmetric positional heterogeneity that tracks with gp41 tilt, which, in turn, correlates with gp120 tilt. Enhanced-sampling simulations showed that FP/FPPR begin to release only at extreme gp120 and gp41 tilt angles. Unbiased simulations of pre-fusion open Env in a viral membrane mimic bound to host membrane-embedded CD4 and CCR5 receptors revealed high gp120 and gp41 tilt angles. Together, these results support a sequential model in which the geometry required for co-receptor engagement forces gp120 into an extreme tilt, propagating to gp41 and pulling the W623 gate away from M530. FPPR release frees the fusion peptide for transit to the host membrane through pre-hairpin formation, and the subsequent HR2 gate releases gp120 (Figure 5E).

Extensive structural data exist for the prefusion closed and receptor CD4-induced open Env trimer and the late pre-hairpin to the post-fusion 6-helix bundle, but the transient structures between the open and late pre-hairpin remain a major gap in our understanding of viral entry. The absence of a clear allosteric rearrangement connecting the co-receptor binding site to the fusion machinery in the CCR5-bound gp120 crystal structure left the question of how the fusion event is triggered. The M530 residue was the clear linchpin holding the FPPR, and therefore the FP, from release, but a plausible trigger for release remained uncertain. The mechanism described here resolves the trigger mechanism without requiring a conventional allosteric path. The signal is geometric, transmitted via trimer tilt imposed by the need to reach a receptor, rather than via classical internal domain rearrangements. Although previous work has suggested that gp120 shedding is not an obligate component of fusion^46^, the fact that disulfide-linked gp120-gp41 virion Envs are fusion-incompetent suggests that eventual gp120 departure is required.^47^ Our results suggest that the timing of gp120 release is critical. Premature shedding would sever the receptor tether that holds the virus at an optimal distance from the host membrane while the FP transits. Placing gp120 release downstream of FPPR release, under control of a second gate, ensures that the trimer retains its host anchor until the fusion peptide has engaged. The same ordering would keep the pre-hairpin transiently shielded and poorly accessible to neutralizing antibodies. Thus, the fusion peptide release and pre-hairpin formation are tightly choreographed events that ensure the process is initiated only when the virus-host system is properly assembled.

The domain tilt-gated system described here likely shifts the probability of M530 release, rather than switching M530 between discretely sequestered and released states. The cryo-EM densities show that the distal alpha-8 W623 gate is poorly resolved, even in the prefusion closed state. This is consistent with positional structural heterogeneity that could permit M530 release. Further, the CD4-induced disassembly mediated by FPPR release suggests that M530 release does occur with Env opening. As observed in the molecular simulations, although rare, a greater gp120 tilt than in the cryo-EM structure does occur, which could lead to premature FPPR release. Thus, the W623 gate is likely coupled to tilt in a probabilistic sense, with a low baseline release in the absence of co-receptor giving way to high-probability release upon the extreme tilt imposed by co-receptor engagement. The relationship between the soluble ectodomain structure and that of the full-length, membrane-embedded Env is also critical. To date, most interpretations have relied on SOSIP to characterize the influence of Env binding to soluble CD4. However, Env, CD4, and CCR5 are transmembrane proteins. Our simulations show that membrane-embedded Env and its association with membrane-embedded CD4 and CCR5 show shifts in gp120 rotation beyond prefusion-stabilized ectodomain open structures. Recent evidence shows that membrane-embedded Env differs from the soluble ectodomain at the gp41 trimer base.^48^ These differences likely affect the clasp and gates and may further reduce premature M530 release, thereby ensuring maximal viable surface Env for host membrane fusion.

Class I viral fusion proteins all transition from metastable trimeric pre-fusion assemblies to alpha-helical coil-coils through pre-hairpin intermediates.^49–51^ In addition to the HIV-1 late pre-hairpin cryo-ET results^2^, extended pre-hairpin intermediates have been captured at resolutions ranging from near-atomic to tomographic envelopes for influenza^52^ and coronaviruses^53,54^. These structural states all depict trimers after the release of their respective fusion-protein peptides and the assembly of the extended pre-hairpin three-helix bundle. The structures that lead to FP release through to pre-hairpin assembly have not been observed. This is likely due to the dynamic and transient nature of these structures. The results here show that this process is tightly regulated in the HIV-1 Env. While pre-hairpin helical bundles and post-fusion coiled coils are shared structural states in class I fusion proteins, the FP release mechanisms likely differ markedly. The receptor binding domains vary widely among class I fusion proteins. Since the FPs are buried in subunit interfaces, the structural states that give rise to release must differ to some degree. Nevertheless, the results here show that FP release is a complex process that gates terminal stages of fusion protein viability. That viability depends on structural features throughout the protein that likely constrain viral evolution beyond coiled-coil formation.

## Methods

### Recombinant HIV-1 Env SOSIP production

A chimeric CH505 transmitted founder (CH505) envelope SOSIP and an N332 BG505 envelope SOSIP^27^ were expressed in Freestyle 293-F cells (Thermo Fisher Scientific, catalog no. R79007). Before transfection, cells were diluted in Freestyle 293 Expression Medium (catalog no. 12338018) to 1.25 × 10^6^ cells/ml at a volume of 950 ml. Plasmid DNAs expressing the envelope SOSIP and furin were co-transfected at a 4:1 ratio (650 and 150 μg per transfection liter, respectively) and incubated with 293fectin transfection reagent (Thermo Fisher Scientific, catalog no. 12347019) in Opti-MEM I Reduced Serum Medium (Thermo Fisher Scientific, catalog no. 31985062) to allow for complex formation. The diluted mixture was added to the cell culture, which was incubated at 37°C, 9% CO2 on a shaker at 120 rpm for 6 days. On day 6, the cell supernatant was harvested by centrifuging the cell culture at 4000*g* for 45 min. The supernatant was filtered with a 0.45-μm polyethersulfone (PES) filter and concentrated to approximately 100 ml using a Vivaflow 200 cross-flow cassette (Sartorius, catalog no. VF20P2). Envelope SOSIPs were purified using a PGT145 affinity chromatography column equilibrated in 15 mM HEPES and 150 mM NaCl (pH 7.1). A PGT145 Ab affinity column was made by coupling PGT145 mAbs to CNBr-activated Sepharose 4B (catalog no. 170430-01, GE Biosciences) and packed into a Tricorn column (GE Healthcare). The supernatant was applied over the column at 2 ml/min using an AKTA go chromatography system (Cytiva) followed by three-column volume wash steps with 15 mM HEPES and 150 mM NaCl. Protein was eluted off the column using 3 M MgCl2 and diluted in 15 mM HEPES and 150 mM NaCl buffer. The protein sample was buffer-exchanged into 15 mM HEPES and 150 mM NaCl by ultrafiltration using a 100-kDa molecular weight cutoff Amicon Ultra-15 Centrifugal Filter Unit (Millipore Aldrich, catalog no. UFC9010) and concentrated to <0.5 ml for size exclusion chromatography. Size exclusion chromatography was performed using a Superose 6 10/300 GL Column (Cytiva) on an AKTA go system in 15 mM HEPES and 150 mM NaCl. Fractions containing trimeric SOSIP were collected. All lots produced were subjected to quality control including analytical size exclusion chromatography, SDS-PAGE, thermal shift analysis, BLI, and NSEM to assure the presence of well-folded Env trimers.

### Recombinant antibody and antibody Fab production

Antibodies were expressed in Expi293F cells in the Expi293 Expression System (Thermo Fisher Scientific, catalog no. A1435101). Cells were diluted to 1.25 × 106 cells/ml at a volume of 100 ml. Heavy-and light-chain plasmid DNAs, at a ratio of 1:1 (50 μg each per 100-ml transfection), were mixed in Opti-MEM I Reduced Serum Medium (Thermo Fisher Scientific, catalog no. 31985062) and incubated with Expifectamine 293 transfection reagent (Thermo Fisher Scientific, catalog no. A14525) to create the DNA-reagent complex. The mixture was added to the prepared cell culture, which was transferred to be incubated at 37°C, 8% CO2 on a shaker at 120 rpm for 6 days. On day 6, the cell supernatant was harvested by centrifuging the cell culture at 4500 rpm for 45 min. The supernatant was filtered with a 0.45-μm PES filter. Each antibody was purified with a Cytiva HiTrap Protein A HP antibody purification column (Cytiva, catalog no. 29048576) equilibrated in 1× phosphate-buffered saline (PBS). The harvested supernatant was applied over the equilibrated column at a flow rate of 0.5 ml/ min using an AKTA start chromatography system (Cytiva), followed by three-column volume wash steps of 1× PBS. The antibodies were then eluted off with Pierce immunoglobulin G (IgG) elution buffer (pH 2.8; Thermo Fisher Scientific, catalog no. 21004). The sample buffer was neutralized with tris (pH 9) at 100 μl/ml (Thermo Fisher Scientific, catalog no. J60707). Lots produced were subjected to quality control including SDS-PAGE and thermal shift analysis. Antibody Fabs were prepared using Lys-C digestion. A digestion mixture of Lys-C enzyme (Thermo Fisher Scientific, catalog no. 90307) and IgG (1:2000 ratio) was combined in 600 μl of 1×PBS for each sample. Each sample was incubated at 37°C for 2 hours. Following the incubation, the digested mAb was removed from the incubator, and a proteinase inhibitor is added to the solution to stop the digestion process. The sample was then purified with a Cytiva HiTrap Protein A HP antibody purification column (Cytiva, catalog no. 29048576), collecting the Fab fragments that flow through. The Fab fragments are then buffer-exchanged into 15 mM HEPES and 150 mM NaCl. Each Fab lot produced was subjected to quality control by SDS-PAGE and thermal shift analysis.

### Differential scanning fluorimetry

Thermal shift assay was performed using Tycho NT.6 (NanoTemper Technologies). Envelope ectodomain SOSIPs were diluted (0.15 mg ml−1) in 15 mM HEPES buffer with 150 mM NaCl at pH 7.1. Intrinsic fluorescence was recorded at 330 and 350 nm while heating the sample from 35° to 95°C at a rate of 3°C min−1. The ratio of fluorescence (350/330 nm) and the inflection temperatures (*T*i) were calculated by Tycho NT.6.

### Biolayer interferometry

Sample mAb and Fab binding were obtained using BLI (OctetR4, ForteBio). Antibodies were immobilized on anti-human IgG Fc capture (ForteBio) sensor tips, while Fabs were immobilized on protein A sensor tips, each via immersion in mAb (4.5 μg/ml) in 15 mM HEPES buffer with 150 mM NaCl at pH 7.1 for 120 s followed by washing in 15 mM HEPES buffer with 150 mM NaCl at pH 7.1 for 60 s at 1000 rpm. The sensor tips were then immersed in the SOSIP-containing wells (45 μg/ml) for 300 s. Reported binding corresponds to values and the end of the association phase. Data were evaluated using the Octet Data Analysis software (ForteBio).

### Negative stain electron microscopy

SOSIP Env samples were incubated with a 5 fold molar excess of either soluble 2 domain CD4 or soluble 2 domain CD4 and 17b antibody Fab for 30 minutes followed by flash freezing in liquid nitrogen. Samples were thawed at room temperature and immediately diluted to 20 μg/ml with 0.1% (w/v) *n*-dodecyl β-d-maltoside [50 mM tris (pH 7.4), 150 mM NaCl, and sodium deoxycholate (0.03 mg/ml)] followed by application to a glow-discharged carbon-coated EM grid for 8 to 10 s, then blotted, and stained with uranyl formate (2 g/dl) for 1 min, blotted, and air-dried. Grids were examined on a Philips EM420 electron microscope operating at 120 kV and nominal magnification of ×49,000, and 114 images were collected on a 76-megapixel charge-coupled device camera at 2.4 Å per pixel. Images were analyzed by 2D class averages using standard protocols with Relion 3.0.^55^

### Cryo-EM sample preparation, data collection, and processing

Purified envelope SOSIP ectodomain preparations were prepared at concentrations of 4 to 5 mg/ml in 15 mM HEPES buffer with 150 mM NaCl at pH 7.1 and mixed with Fab at a 1:5 molar ratio. A total of 2.5 μl of the complex was deposited on a CF-1.2/1.3 grid that had been glow-discharged for 15 s in a PELCO easiGlow glow-discharge cleaning system. After 30-s incubation in >95% humidity, the excess protein was blotted away for 2.5 s before being plunge-frozen in liquid ethane using a Vitrobot Mark IV plunge freezer (ThermoFisher Scientific). Frozen grids were imaged in a Talos Arctica (ThermoFisher Scientific) equipped with a K3 detector (Gatan). Individual frames were aligned, dose-weighted, and contrast transfer function (CTF) corrected, followed by particle picking, 2D classification, ab initio model generation, heterogeneous refinements, homogeneous 3D refinements, and local resolution calculations in cryoSPARC.^56^ Local refinement of the disordered FPPR/alpha-8 protomer was performed for the CH505 parent and D624P construct close-state maps. Masks for particle subtraction and refinement were obtained using ChimeraX^57^ to segment maps into single protomers. The resulting maps were used to evaluate FPPR and alpha-8 secondary structures and in final structure refinements to fit secondary structure, followed by final fitting to the full trimer maps.

### Cryo-EM structure fitting and analysis

The CH505.M5.G458Y SOSIP structure (6UDA) and CH235.12 structures were used to fit the cryo-EM maps in ChimeraX.^57^ Mutations were made in PyMol.^58^ Coordinates were then fitted using Isolde^59^ followed by iterative refinement using Phenix^60^ real-space refinement. Structure and map analyses were performed using PyMol and ChimeraX. Protomer angular terms for gp120 and gp41 were calculated using W571 and W596 c-αs, and the c-αs of gp120 centroid, excluding variable loops, the V1/V1 region residues, and the c-αs of residues W589 and W631 of the protomer and residue W589 of the adjacent protomer (clockwise viewing from the trimer apex), respectively.

### Cryo-EM Volume Density Calculation

Cryo-EM maps were normalized to standardize their density distributions between 0 and 1 and displayed at a common contour level of 0.5 for direct comparison. Map volumes were quantified within regions of interest defined by a 3 Å radius around the selected atoms. Measurements were performed for the full protomer, including gp120 and gp41, the fusion peptide–proximal region (FPPR; residues 530–540), and the α8 region (residues 610–628). Quantified volumes are reported in Å³. For each trimer, the protomer with the highest measured map volume was designated protomer 1 and assigned a relative value of 1.0. The remaining protomers were ranked in descending order and relative to protomer 1. Only reconstructions with global resolutions of 4.5 Å or better were included in the quantitative analysis. CH505 pre-fusion open Env reconstructions with global resolutions poorer than 4.5 Å showed insufficient local map quality around the atomic models and were therefore excluded. All map processing and volume measurements were performed in ChimeraX, and the data were plotted in GraphPad Prism.

### gp120/41 base molecular dynamics simulations

The gp120/gp41 base system was prepared using the CD4 and 17b-bound open-state B41 isolate (PDB ID 5VN3^8^) structure. Residues 32-47 and 489-505 of gp120 and 584-653 of gp41 were retained, and the 47 and 489 backbone residues of gp120 were linked to form a single gp120-terminus peptide chain. The resulting structure was minimized using the Schrödinger^61^ molecular modeling package. The protein was immersed in a truncated-octahedral TIP3P water box with a minimum edge distance of 12 Å to the nearest protein atom. A chlorine atom was added to neutralize the system charge, and ionizable residue protonation states were assigned based on a pH of 7.0. The Amber ff14SB protein^62^ force fields were used throughout. All simulations were performed using the Amber20 pmemd CUDA implementation. The systems were first minimized for 10,000 steps with protein atom restraints, followed by minimization of the full system without restraints for an additional 10,000 steps. This was followed by heating the systems from 0 K to 298 K over 20 ps in the NVT ensemble using a 2-fs time step and the particle-mesh Ewald method for long-range electrostatics, with periodic boundary conditions.^63^ The systems were then equilibrated for 100 ps in the NPT ensemble, with temperature controlled by Langevin dynamics at 1.0 ps^-1^ and 1 atm pressure maintained by isotropic position scaling with a relaxation time of 2 ps.^64^ A non-bonded cutoff of 9 Å was used throughout and hydrogen atoms were constrained using the SHAKE algorithm^65^ with hydrogen mass repartitioning^66^ used to allow for a 4-fs time step. A total of 25 independent, 5 microsecond production simulations were conducted using a weak, 1.0 kcal/mol/Å^2^ harmonic restraint on HR1 three-helix bundle helix residues 584-595 to mimic the trimeric context.

### Functional assay cells, plasmids, viruses

HEK293T (ATCC CRL-3216), HeLa TZM-bl ^67^ (NIH AIDS Reagent program catalog no. 8129), and HEK293T-huCD4/CCR5 ^68^, which are engineered to express human CD4 and CCR5, cells were cultured in Dulbecco’s modified eagle medium (DMEM, Gibco) supplemented with 10% fetal bovine serum (FBS, Sigma), 2 mM L-glutamine (Gibco), and 1x Penicillin-Streptomycin (Gibco) (complete DMEM). Full-length, replication-competent HIV-1 Q23 BG505 infectious molecular clone (HIV-1 BG505) was used to construct mutant Env variants. Briefly, the MfeI–SalI region of BG505.W6M.B1 Env encoding the D624P, N637P, and D624P+N637P mutations was synthesized as DNA fragments (Twist Bioscience). HIV-1 BG505 D624P, HIV-1 BG505 N637P, and HIV-1 BG505 D624P+N637P proviral clones were generated by digestion and ligation of these fragments into the HIV-1 BG505 proviral plasmid using MfeI and SalI restriction sites. All resulting proviral plasmids were verified by Sanger DNA sequencing. β-lactamase (BlaM) gene fused to the N-terminus of HIV-1 Vpr (Genbank: AF324493) separated by six-glycine-one-lysine linker was synthesized as a DNA fragment (Integrated DNA Technologies), digested, and ligated into pcDNA3.1 (Invitrogen) using KpnI and NotI restriction sites to generate pcDNA3.1-Blam-HIV-1 Vpr plasmid. The resulting plasmids were verified by Sanger DNA sequencing. Replication-competent HIV-1 were generated by transfecting 2 x 10^6^ HEK293T cells with 4 μg of proviral plasmid DNA and 12 μl of Fugene 6 transfection reagent (Roche) following manufacturer’s protocol. Forty-eight hours post-transfection, virus-containing supernatant was harvested, passed through a 0.2 μm sterile filter and concentrated ∼10-fold using Amicon Ultracel 100 kDa filters (Millipore).

### TZM-bl infectivity assay

10-fold serial dilutions (10^-1^ to 10^-6^) of HIV-1 stocks were added to 2 x 10^4^ TZM-bl cells in the presence of 10 μg/ml DEAE-dextran. Forty-eight hours post-infection, cells were fixed in 500 μl of fixing solution (1% formaldehyde and 0.2% glutaraldehyde in 1x PBS) for 5 minutes, washed twice with 1x PBS, stained for β-galactosidase for 50 minutes at 37°C in 200 μl of staining solution (0.4 mg/mL X-gal, 2 mM MgCl_2_, 4 mM K_3_[Fe(CN)_6_], 4 mM K_4_[Fe(CN)_6_] in 1x PBS), washed twice with 1x PBS, and blue foci were counted to quantify HIV-1 titers.

### Reverse transcriptase activity assay

RT activity assay was performed as described previously ^69^. Briefly, 5 µl of HIV-1 stock or RT standard was lysed in 5 µl of 2x lysis buffer (100 mM Tris HCl pH 7.4, 50 mM KCl, 0.25% Triton X-100, 40% glycerol) in the presence of 4U RNaseOUT (Invitrogen) for 10 minutes at room temperature. Viral lysate was diluted 1:10 by adding 90 µl of nuclease-free water (Life Technologies). qRT-PCR reactions were prepared by mixing 9.6 µl of diluted viral lysate with 10.4 µl of reaction mix containing 10 µl of 2x Maxima SYBR Green/ROX qPCR Master Mix (ThermoFisher), 0.1 µl of 4U/µl RNaseOUT, 0.1 µl of 0.8 µg/µl MS2 RNA template (Roche), and 0.1 µl each of 100 µM forward 5’-TCCTGCTCAACTTCCTGTCGAG-3’ and reverse 5’-CACAGGTCAAACCTCCTAGGAATG-3’ primers. qRT-PCR was performed using a QuantStudio 3 Real-Time PCR machine (Applied Biosystems). Viral titers were calculated from a standard curve generated using recombinant reverse transcriptase (Millipore catalog no. 382129).

### HIV-1 BlaM-Vpr virus fusion assay

The HIV-1 β-lactamase-Vpr (BlaM-Vpr) virus fusion assay was performed as described previously ^70,71^ with some modifications. Briefly, HIV-1 containing Blam-Vpr fusion protein were generated by co-transfecting HEK293T cells with 4.5 μg of proviral HIV-1 plasmid and 1.5 µg of pcDNA3.1-Blam-Vpr plasmid using Fugene 6 transfection reagent following manufacturer’s protocol. Forty-eight hours post-transfection, virus-containing supernatant was harvested, passed through a 0.2 μm sterile filter, concentrated ∼10-fold using Amicon Ultracel 100 kDa filters (Millipore), aliquoted and stored at-80°C. RT activity of viral stocks was measured using the RT activity assay. 3 x 10^5^ HEK293T-huCD4/CCR5 cells in 100 µl of complete DMEM were infected with Blam-Vpr-containing HIV-1 equivalent to 350 mU of RT by spinoculation at 1200 x g for 90 minutes followed by incubation at 37°C and 5% CO_2_ for one hour. Fusion-mediated HIV-1 entry was quantified by monitoring the conversion of fluorescent BlaM CCF2-AM substrate dye as described. After infection, cells were washed once with 300 µl of cold CO_2_-independent media (Gibco) without FBS and resuspended in 100 µl of CO_2_-independent media supplemented with 10% FBS. Cells were incubated with CCF2-AM substrate (LiveBLAzer CCF2-AM Kit, Invitrogen) following manufacturer’s protocol in the presence of 1.8 mM Probenecid (Sigma) for 2 hours. Cells were washed three times with 300 µl of cold CO_2_-independent media without FBS, once with 1x PBS, fixed with 200 µl of 2% paraformaldehyde, washed once with 1x PBS, and resuspended in 400 µl of FACS buffer. The fluorescence of cleaved CCF2-AM substrate was measured on CytoFlex S flow cytometer and data were analyzed using FlowJo software.

### Env shedding assay

HEK293T cells (2 x 10^6^) were transfected with 4 μg of proviral HIV-1 plasmid using Fugene 6 transfection reagent following manufacturer’s protocol. Virus-containing supernatants were collected at 6, 12, 24, 36, 48, and 60 hours post-transfection, clarified by filtration through a 0.2 μm filter, and pelleted through a 25% sucrose cushion by ultracentrifugation at 28,000 rpm for 90 minutes at 4°C. Post-ultracentrifugation supernatants were collected for analysis. For detection of shed Env, 36 μl of post-ultracentrifugation supernatant was mixed with 12 μl of 4× Laemmli sample buffer (BioRad) and 2 μl of 2-mercaptoethanol, then subjected to SDS-PAGE followed by standard Western blotting. Env levels were detected using HIV-1 anti-gp120 antibody (NIH AIDS Reagent Program, catalog no. 288). Band intensities corresponding to gp160 and gp120 were quantified using ImageJ software.

### Immunoblotting

Whole cell extracts were prepared by lysing the cells in radioimmunoprecipitation assay (RIPA) cell lysis buffer (50 mM Tris pH 8.0, 0.1% SDS, 1% Triton-X, 150 mM NaCl, 1% deoxycholic acid, 2 mM PMSF). For virion incorporation, virus containing supernatants from the infected cell cultures were centrifuged at 650 x g for five minutes at room temperature. Cell-free supernatant was filtered through 0.2 µm filter and then pelleted through a 25% sucrose cushion by ultracentrifugation for at 28,000 rpm for 90 minutes at 4°C. Virus pellets were lysed in 70 μl of RIPA buffer for 10 minutes at room temperature. The lysates were normalized by RT activity, subjected to SDS-PAGE and immunoblotted. Standard Western blotting procedures were used with the following antibodies: HIV-1 gp120 (NIH AIDS Reagent program catalog no. 288), HIV-1 p24 (NIH AIDS Reagent program catalog no. 6521), and β-Actin (Proteintech catalog no. 66009-1-Ig). Protein expression was quantified by measuring the band intensities of Env and p24 using ImageJ software.

### Gaussian accelerated molecular dynamics and ligand Gaussian accelerated molecular dynamics 3 simulations of the open SOSIP trimer

We performed GaMD and LiGaMD3 simulations on the open SOSIP trimer to explore the sequential conformational transitions required for the FP to emerge from its pocket for fusogenic competence. The simulation system was prepared starting from the 5VN3^72^ PDB structure, the cryo-EM model of B341 SOSIP.664 in complex with soluble CD4, with only the D1 and D2 domains, and antigen binding variable domain of the 17b antibodies. Since we did not model the full CD4 receptors in the T-cell membrane in this study to simulate its pulling effects on the gp120, we removed the soluble CD4 and the 17b antibodies from the structure. We then used MODELLER to model the complete structure of the open B41 SOSIP based on the gp120 and gp41 sequences included in Supplementary Table 1. We capped the N-and C-termini of the gp120 and gp41 subunits with neutral patches, including acetyl and methylamide, respectively. The modelled open B41 SOSIP was then embedded in 0.15 M NaCl solution using the CHARMM-GUI webserver ^73,74^, resulting in a simulation system box dimension of 184.68 Å × 184.68 Å × 184.68 Å and simulation system size of 627,357 atoms.

The CHARMM36m force field parameter set ^75^ was employed to parameterize the simulations. The standard builds of the simulation software AMBER 24 ^76^ were applied to carry out the simulations, using both the CPU and GPU version of the *pmemd* module (*pmemd* and *pmemd.cuda*). We used the simulation input files prepared by the CHARMM-GUI webserver ^73,74^ to perform energetic minimization, equilibration with the constant number, volume, and temperature (NVT) ensemble, and a short 25-ns conventional MD (cMD) simulation using the constant number, pressure, and temperature (NPT) ensemble to equilibrate the simulation system prior to the GaMD^44^ and LiGaMD3^45^ simulations. We applied the periodic boundary conditions and restrained the bonds containing hydrogen atoms using the SHAKE^65^ algorithm. We kept the temperature constant at 310 K using the Langevin thermostat^77,78^ using a friction coefficient of 1.0 ps^-1^. We calculated the electrostatic interactions using the particle mesh Ewald summation^79^ with a cutoff of 9.0 Å for long-range interactions. We kept the pressure constant at 1.0 bar using the Berendsen barostat^80^ with isotropic coupling. The initial energetic minimization of the simulation system was carried out using the steepest-descent algorithm. We used a timestep of 2 fs for the cMD simulation and the following GaMD^44^ and LiGaMD3^45^ simulations. In the all-atom dual-boost GaMD^44^ simulations, one boost potential was applied to the total potential energetic term, while another boost potential was applied to the dihedral energetic term of the simulation system (Supplementary Table 2). The reference energy *E* was set to a lower bound (*E = V_max_* ^44^) for both the total and dihedral potential energy (Supplementary Table 2). We set the upper limits of the boost potential standard deviations for both the total potential and dihedral energetic terms, σ_0P_ and σ_0D_, to 6.0 kcal/mol (Supplementary Table 2). In the selective LiGaMD3^45^ simulations, one boost potential was added to the essential nonbonded potential energy of the first 11 residues of the fusion peptides (including A512 – G522), another boost potential was added to the remaining nonbonded potential energy of the whole system, and the last boost potential was added to the total bonded potential energy of the system (Supplementary Table 2). The first boost potential was only added to the first 11 residues of the fusion peptides due to the constraints for the maximum number of atoms in the TI mask of 500 atoms in AMBER^76^. The first and second reference energy corresponding to the first and second boost potentials were set to upper bounds (*E = V_min_ + (V_max_ –V_min_) / k_0_*^45,81^), while the third reference energy was set to a lower bound (*E = V_max_*^44,45^) (Supplementary Table 2). The upper limits of the boost potential standard deviations for the first, second, and third boost potentials (σ_0P_, σ_0D_, and σ_0B_, respectively) were set to 7.0 kcal/mol, 6.0 kcal/mol, and 6.0 kcal/mol (Supplementary Table 2). Both the all-atom dual-boost GaMD^44^ and LiGaMD3^45^ simulations of the open B41 SOSIP trimer first involved an initial cMD simulation of 25 ns to calculate the acceleration parameters, followed by the equilibration stage of 100 ns for the initial addition of boost potential and update of the acceleration parameters. In total, the GaMD^44^ and LiGaMD3^45^ equilibration simulations each lasted for 125 ns. Three independent 1000 ns GaMD^44^ production simulations and three independent 1250 ns LiGaMD3^45^ production simulations were carried out with randomized initial velocities using the acceleration parameters calculated from the GaMD^44^ and LiGaMD3^45^ equilibration simulations.

### Reaction coordinates (RCs) to monitor FP/FPPR dynamics

Following the production simulations, we employed the CPPTRAJ^82^ module within the AmberTools24^83^ to analyze the simulation trajectories. We used the HXB2 numbering scheme for gp160 to assign residue numbers within the HIV-1 B41 SOSIP. In particular, we calculated the time courses of the experimentally recommended Reaction coordinates (RCs), including the C_α_-atom distances between gp41 residue 571 and the gp120 COM (including residues 32-129 and 198-395) (namely, distance 1), C_α_-atom distances between gp41 residues 571 and fusion peptide proximal region (FPPR) center-of-mass (COM) (consisting of gp41 residues 528-540) (distance 2), C_α_-atom distances between gp41 residues 530 (of the FPPR) and 623 (of the W-clasp) (distance 3), dihedral angle (defined by the gp120 COM, C_α_-atom COM of gp120 residues 46 and 396, C_α_-atom of gp41 residue W596, and C_α_-atom of gp41 residue W571) (namely dihedral angle), angle 1 (defined by the gp120 COM, C_α_-atom COM of gp120 residues 46 and 396, and C_α_-atom of gp41 residue 596), angle 2 (defined by the C_α_-atom COM of gp120 residues 46 and 396, C_α_-atom of gp41 residue 596, and C_α_-atom of gp41 residue 571), gp120 tilt angle (or gp120 angle, defined by C_α_ atoms of gp120 COM and the center z-axis drew through the gp41 three-helix bundle), and gp41 tilt angle (or gp41 angle, defined by C_α_ atoms of gp41 residues 566-596 COM and the center z-axis drew through the gp41 three-helix bundle). Afterward, we plotted two-dimensional (2D) Gaussian kernel density estimation (KDE) from pairs of calculated RCs to identify the conformational states sampled from the GaMD and LiGaMD3 simulations.

### All-atom MD simulations of fully open Env in complex with CD4 and CCR5 in complex membrane

We assembled existing Cryo-EM structures to generate the initial complex structure of our simulations. The fully open Env structure was obtained from PDB ID: 5VN3^72^, with 17b removed. The missing loops in 5VN3 were modelled using SWISS-MODEL^84^. The interface between CD4 and gp120 was modelled based on PDB ID: 5VN3^72^. The interface between CCR5 and gp120 was modelled based on PDB ID: 6MET^25^. The D1-D4 domain of CD4 was modelled from PDB ID: 6MET^25^ and the transmembrane domain of CD4 was modelled from PDB ID: 2KLU^85^ using SWISS-MODEL^84^. For modeling the full length gp41 structure, the ectodomain was obtained from PDB ID: 5VN3^72^ and the CT, TM and MPER region were obtained from PDB ID: 7LOI^86^. We used SWISS-MODEL^84^ to connect the gp41 ectodomain structure to the MPER region. The TM domain of gp41 was embedded in viral membrane mimic and the lipid composition was the same as our previous MD simulation study^87^. The TM domain of CD4 and CCR5 were embedded in the T-cell membrane mimic, and the lipid composition was the same as our previous MD simulation study^88^. Embedding of protein into the complex membranes were performed using CHARMM-GUI^74^. The distance between the outer leaflet of T-cell membrane and the outer leaflet of HIV-1 membrane is ∼12 nm, which is shorter than Env in complex with three CD4 bound and is consistent with the speculation that binding to CCR5 bring the host and viral membrane closer together^24^. The sequence of Env used in this study was retrieved from GenBank sequence ID: AGG24726.1 which was the Env sequence from patient CH505^89^. The simulation system was initially put in a triclinic box with dimensions 32.471 x 32.471 x 35 nm and was solvated in 898,763 TIP3P^90^ water molecules with 3702 Na^+^ and 3334 Cl^-^ counter ions to create a 0.15 mol/L salt concentration in the simulation box. There are 3,597,519 atoms in total in the simulation box. The counter ions were placed such that the net charges between the T-cell and HIV-1 membranes are the same as the net charges outside of the membranes. For these MD simulations, we used CHARMM36-LJPME force field released in July 2022 (downloaded from https://mackerell.umaryland.edu/charmm_ff.shtml#gromacs) for protein^75^ (CHARMM36m for protein) and lipids^91^. GROMACS 2025.3^92^ simulation software was used for all the equilibration and simulations. The simulation system was energy minimized using steepest descent algorithm until the maximum force is below 1000 kJ mol^-1^ nm^-1^. The energy minimization was followed by a series of equilibrations in NPT condition with C-rescale pressure coupling, semiisotropic coupling type, time constant = 8.0 ps and compressibility = 4.5E-5 bar^-1^. The following simulations were performed in sequence: 1) NPT equilibration with reference pressure to 10 bar for 1 ns and position restraint on protein with force constant 1000 kJ mol^-1^ nm^-2^, 2) NPT equilibration with reference pressure to 1 bar for 2 ns and position restraint on protein with force constant 1000 kJ mol^-1^ nm^-2^. This step was followed by 4 NPT equilibrations with reference pressure to 1 bar for 1 ns each while reducing the force constant for the position restraint on the protein to 800, 600, 400, 200, kJ mol^-1^ nm^-2^, sequentially. Finally, we performed a short MD simulation in NPT condition without position restraint to the protein for 1 ns. We then performed the four independent production MD simulations for 100 ns each. The long-range electrostatic interactions were calculated using Particle Mesh Ewald algorithm. The short-range coulomb, Van der Waals and neighbor list cut-off distance are 1.2 nm. Temperature coupling was performed using V-rescale with reference temperature of 310 K and time constant of 1 ps.

The gp41 and gp120 tilt (θ angle) angles were calculated for the last half of the simulations. Gp41 tilt angle is defined as the angle between three points, i) C-α of residue 631 of the current protomer being calculated, ii) C-α of residue 589 of the current protomer, iii) C-α of residue 589 of the neighboring protomer in the clockwise direction of the current protomer. Gp120 tilt angle is defined as the angle between three points, i) center of mass of the C-α atoms of Gp120 residues (excluding V1, V2, V4 and V5 loops), ii) center of mass of C-α atoms from residue 46 and 490, iii) C-α of residue 596.

## Acknowledgements

The cryo-EM experiments were carried out at the University of North Carolina cryo-EM Core Facility. We would like to thank Los Alamos National Laboratory Institutional Computing for providing the computing resources to run atomistic simulations. We thank Josh Strauss for technical assistance in this project. We thank Cesar A. López for helpful discussions on setting up the fully open Env in complex with CD4 and CCR5 in a complex membrane simulation system. The cryo-EM facilities were supported by the National Cancer Institute of the National Institutes of Health under award number P30CA016086. This project was supported by the NIH, National Institute of Allergy and Infectious Diseases, Division of AIDS Consortia for HIV/AIDS Vaccine Development (CHAVD) grants UM1AI144371 (to B.F.H.), R01AI145687 (P.A.), and R01AI172615 (A.S.), and U54AI170752 (R.H., S.G and P.A.), and the Translating Duke Health Initiative (R.H. and P.A.).

## Author Contributions

H.D., I.U. and S.G. conducted and analyzed the molecular dynamics simulations; M.L. and P.A. analyzed cryo-EM-derived molecular volumes; D.F., N.R., and A.S. conducted and analyzed functional experiments; A.J. produced and analyzed virus-like particles; A.J. and K.J. obtained and analyzed virus-like particle cryo-EM micrographs; C.S. produced and characterized prefusion-stabilized Env trimer ectodomains; H.L. and R.J.E. obtained and analyzed the negative stain electron microscopy data; R.H. designed trimer gate 1 and gate 2 perturbation mutations, obtained and processed cryo-EM data, and analyzed trimer geometry; All authors participated in the preparation and approved of the manuscript.

## Data Availability

All data presented in this manuscript will be made available in public repositories upon formal publication.

---

Cryo-EM data collection, refinement and validation statistics

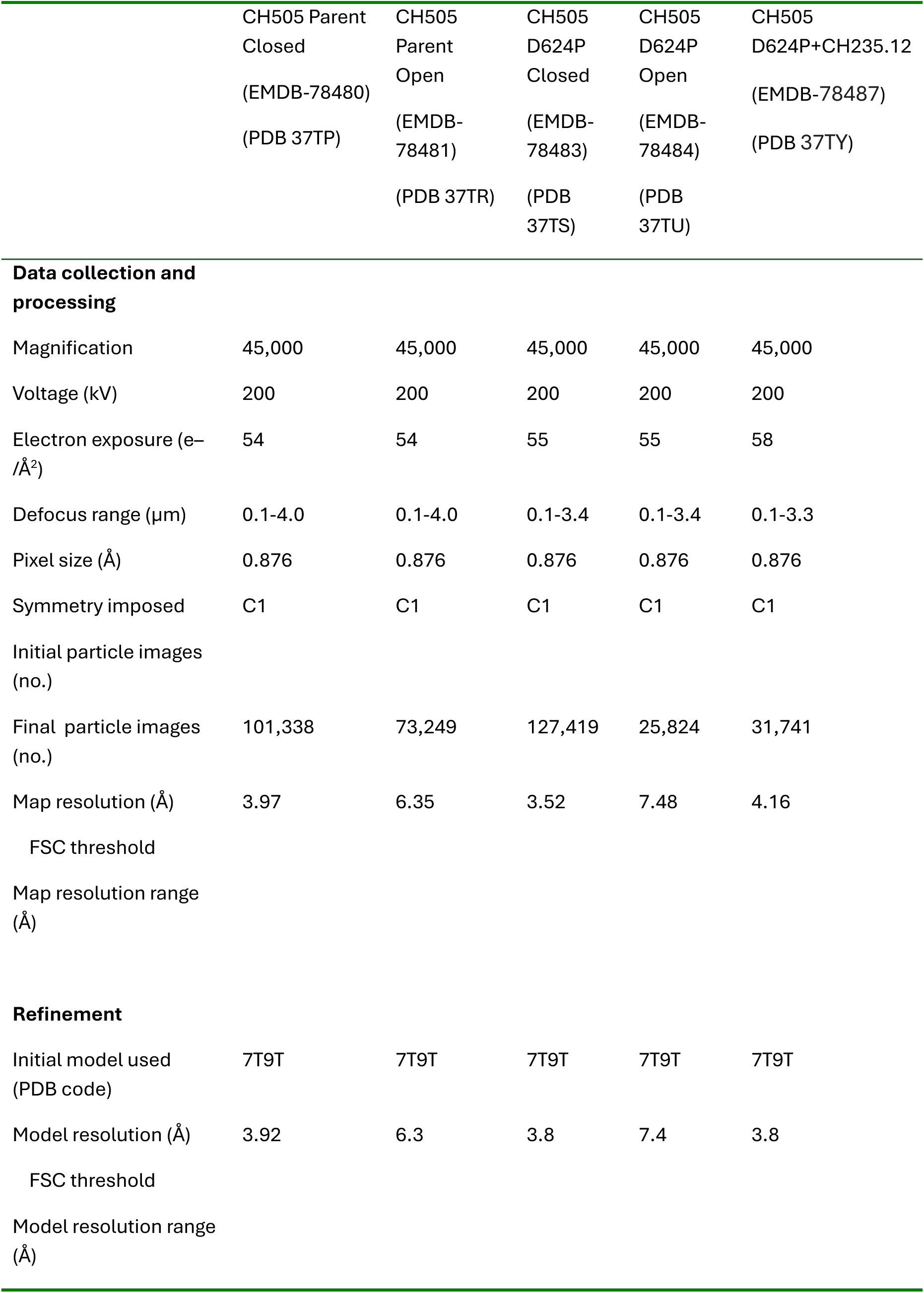
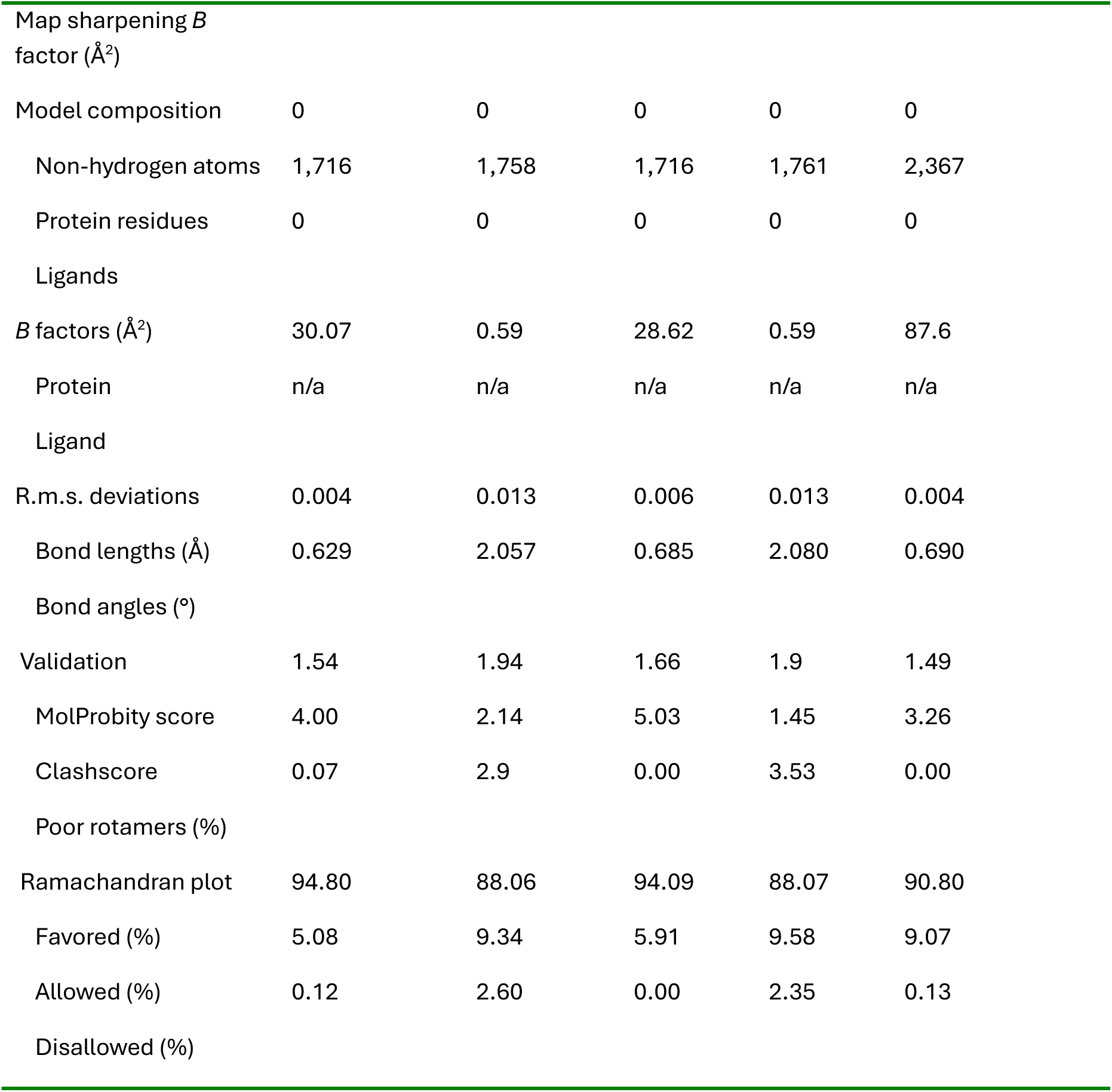

|  |  |  |  |  |  |
| --- | --- | --- | --- | --- | --- |
|  | CH505 Parent<br>Closed<br>(EMDB-78480)<br>(PDB 37TP) | CH505<br>Parent<br>Open<br>(EMDB-78481)<br>(PDB 37TR) | CH505<br>D624P<br>Closed<br>(EMDB-78483)<br>(PDB 37TS) | CH505<br>D624P<br>Open<br>(EMDB-78484)<br>(PDB 37TU) | CH505<br>D624P+CH235.12<br>(EMDB-78487)<br>(PDB 37TY) |
| **Data collection and processing** |  |  |  |  |  |
| Magnification | 45,000 | 45,000 | 45,000 | 45,000 | 45,000 |
| Voltage (kV) | 200 | 200 | 200 | 200 | 200 |
| Electron exposure (e<sup>-</sup>/Å<sup>2</sup>) | 54 | 54 | 55 | 55 | 58 |
| Defocus range (µm) | 0.1-4.0 | 0.1-4.0 | 0.1-3.4 | 0.1-3.4 | 0.1-3.3 |
| Pixel size (Å) | 0.876 | 0.876 | 0.876 | 0.876 | 0.876 |
| Symmetry imposed | C1 | C1 | C1 | C1 | C1 |
| Initial particle images (no.) |  |  |  |  |  |
| Final particle images (no.) | 101,338 | 73,249 | 127,419 | 25,824 | 31,741 |
| Map resolution (Å) | 3.97 | 6.35 | 3.52 | 7.48 | 4.16 |
| FSC threshold |  |  |  |  |  |
| Map resolution range (Å) |  |  |  |  |  |
| **Refinement** |  |  |  |  |  |
| Initial model used (PDB code) | 7T9T | 7T9T | 7T9T | 7T9T | 7T9T |
| Model resolution (Å) | 3.92 | 6.3 | 3.8 | 7.4 | 3.8 |
| FSC threshold |  |  |  |  |  |
| Model resolution range (Å) |  |  |  |  |  |
| Map sharpening *B* factor (Å<sup>2</sup>) |  |  |  |  |  |
| Model composition | 0 | 0 | 0 | 0 | 0 |
| Non-hydrogen atoms | 1,716 | 1,758 | 1,716 | 1,761 | 2,367 |
| Protein residues | 0 | 0 | 0 | 0 | 0 |
| Ligands |  |  |  |  |  |
| *B* factors (Å<sup>2</sup>) | 30.07 | 0.59 | 28.62 | 0.59 | 87.6 |
| Protein | n/a | n/a | n/a | n/a | n/a |
| Ligand |  |  |  |  |  |
| R.m.s. deviations | 0.004 | 0.013 | 0.006 | 0.013 | 0.004 |
| Bond lengths (Å) | 0.629 | 2.057 | 0.685 | 2.080 | 0.690 |
| Bond angles (°) |  |  |  |  |  |
| Validation | 1.54 | 1.94 | 1.66 | 1.9 | 1.49 |
| MolProbity score | 4.00 | 2.14 | 5.03 | 1.45 | 3.26 |
| Clashscore | 0.07 | 2.9 | 0.00 | 3.53 | 0.00 |
| Poor rotamers (%) |  |  |  |  |  |
| Ramachandran plot | 94.80 | 88.06 | 94.09 | 88.07 | 90.80 |
| Favored (%) | 5.08 | 9.34 | 5.91 | 9.58 | 9.07 |
| Allowed (%) | 0.12 | 2.60 | 0.00 | 2.35 | 0.13 |
| Disallowed (%) |  |  |  |  |  |

