## Supplemental Material for "Sequential HIV-1 fusion protein gates control the initiation of host cell entry"

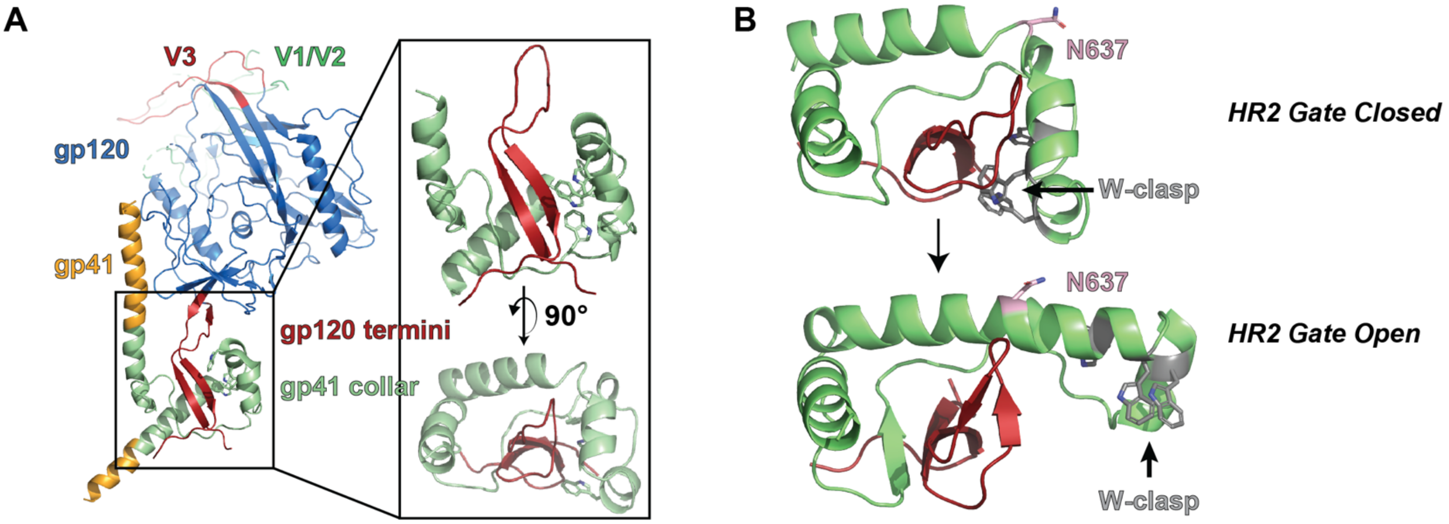


**Supplemental Figure 1. Molecular dynamics simulation of the FP/FPPR free open state gp120-termini/gp41-collar reveals helical extension in HR2 at position N637. (A)** Open state (PDB ID 5VN3) protomer highlighting the simulated system, including the gp120-termini (dark red) and gp41-collar (light green) without the FP/FPPR residues. **(B)** (top) The initial open occluded state conformation highlighting the W-clasp and alpha-8-turn-alpha-8 helical motif containing N637 in the turn. (bottom) A representative structure of the HR2 helical extension.

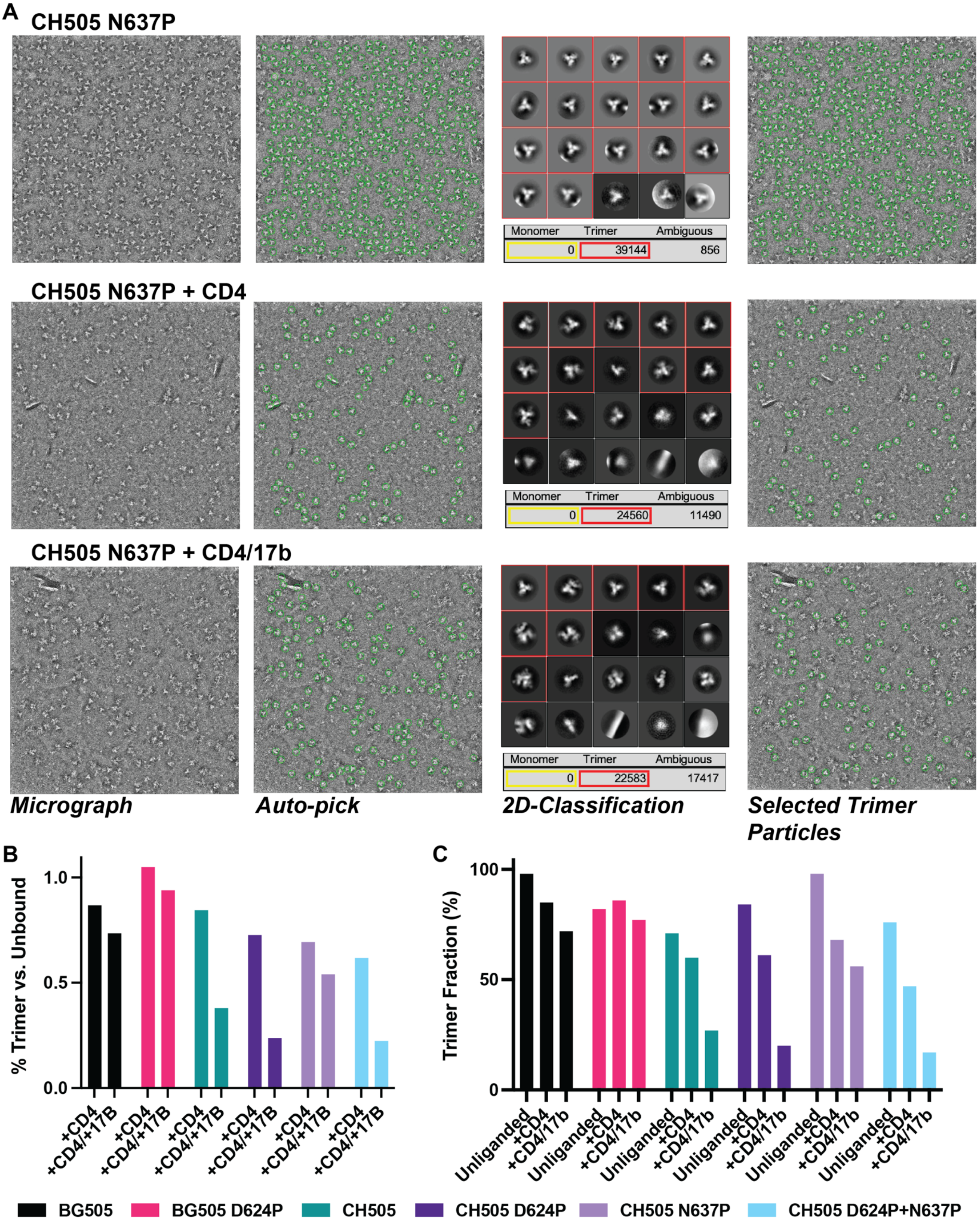


**Supplemental Figure 2. Negative stain electron microscopy Env SOSIP trimer disassembly assay reveals differential impacts of CD4 and 17b interaction on the trimer conformation. (A)** (top-bottom) Representative assay processing steps for the CH505 N637P construct. (left-right) Representative NSEM micrographs, auto-pick selections, 2D-class average images and particle distributions into monomeric, trimeric, and ambiguous classes, final selected trimer particles. **(B)** The +CD4 and +CD4/17b NSEM percent trimer fraction vs. the unbound trimer NSEM. **(C)** Percent trimer fractions for each condition.

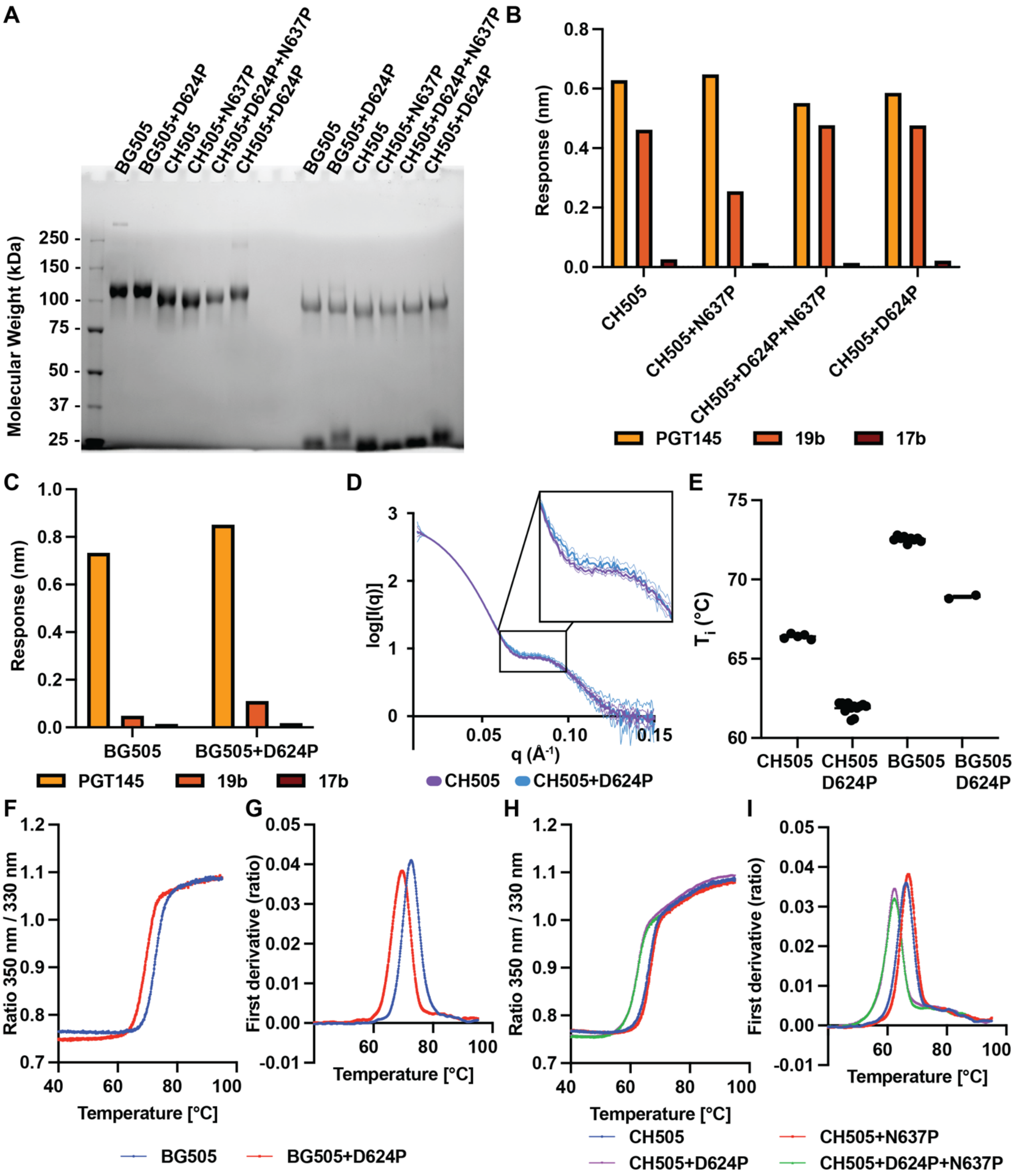


**Supplemental Figure 3. Construct purity, conformation, and stability. (A)** Reducing and non-reducing SDS-PAGE. **(B)** Biolayer interferometry binding data for the CH505 Env SOSIP constructs interacting with the trimer apex targeting PGT145, V3-loop targeting 19b, and bridging-sheet targeting 17b antibodies. **(C)** Biolayer interferometry binding data for the BG505 Env SOSIP constructs interacting with the trimer apex targeting PGT145, V3-loop targeting 19b, and bridging-sheet targeting 17b antibodies. **(D)** Small-angle X-ray scattering curves for the CH505 parent and CH505 D624P constructs highlighting a profile feature that correlates with Env protomer gp120 rotation. (E) Multiple-lot $T_i$ measures for the CH505 (N=5 lots), CH505 D624P (N=12 lots), BG505 (N=9 lots), and BG505 D624P (N=2 lots). One lot represents an independent construct transfection and purification. **(F)** BG505 constructs Tycho DSF denaturation profiles for the absorption ratio between 330 nm and 350 nm during non-equilibrium heating. **(G)** The first derivative profiles of the BG505 ratio plots. **(H)** CH505 construct Tycho DSF denaturation profiles for the absorption ratio between 350 nm and 330 nm during non-equilibrium heating. **(I)** The first derivative profiles of the CH505 ratio plots.

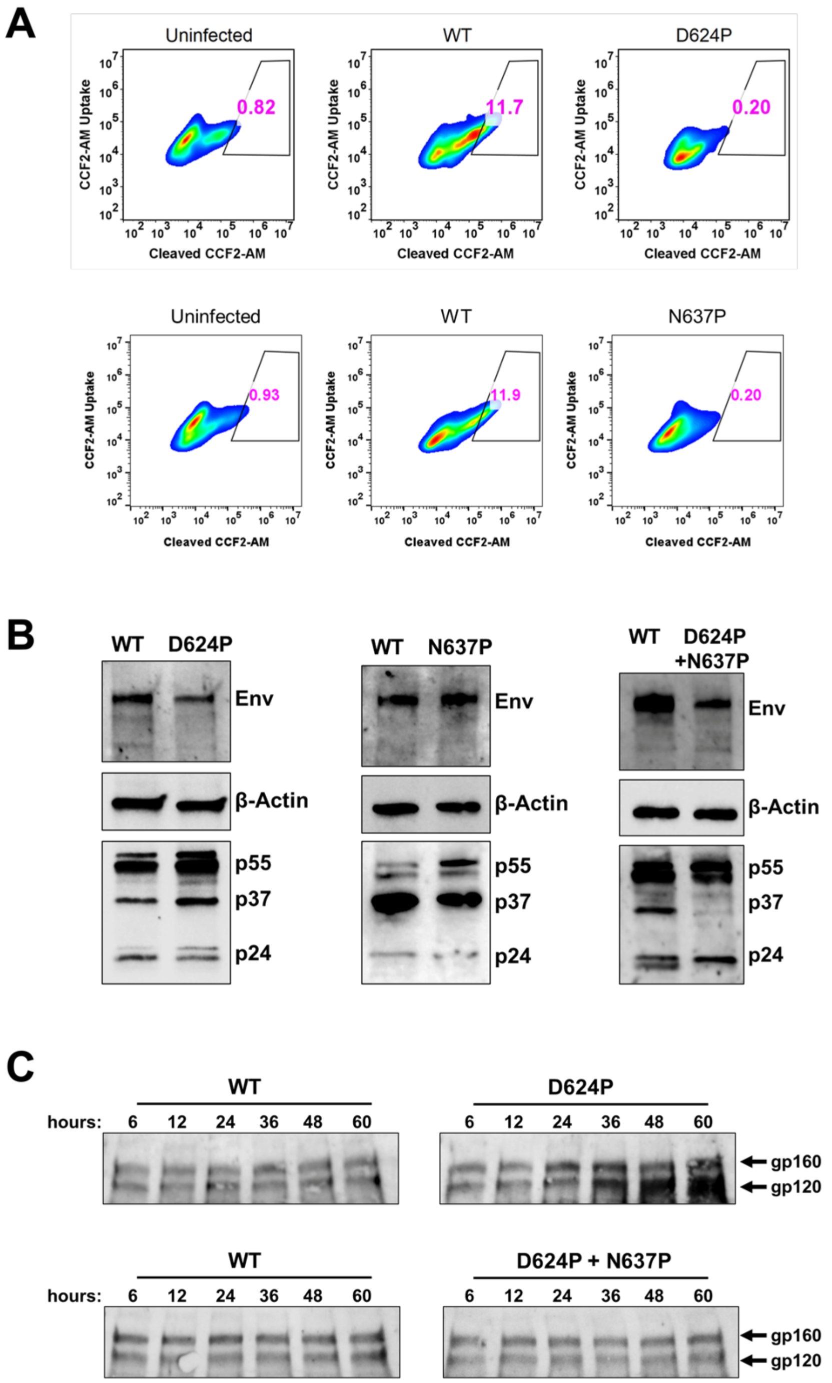


**Supplemental Figure 4.** ***Blocking HR2 helical extension rescues the wild-type shedding phenotype from induced W623 gate instability.*** **A)** Representative flow cytometry plots showing viral fusion, expressed as the percentage of cells positive for cleaved CCF-AM substrate, measured using the HIV-1 BlaM-Vpr fusion assay. Fusion of the indicated HIV-1 variants with HEK293T cells expressing human CD4 and CCR5 is shown, with the Env variant encoded by each virus indicated above the plots. **B)** Western blot analysis of whole cell extracts harvested at 48 hours post-transfection from HEK293T cells transfected with the indicated HIV-1 proviral plasmids. Immunoblotting performed using anti-Env, anti-p24, and anti-actin antibodies. **C)** Representative Western blot analysis of gp120 shedding over time. Post-ultracentrifugation supernatants collected at the indicated time points (6, 12, 24, 36, 48, and 60 hours) from HEK293T cells transfected with the indicated HIV-1 proviral plasmids. Immunoblotting performed using an anti-Env antibody.

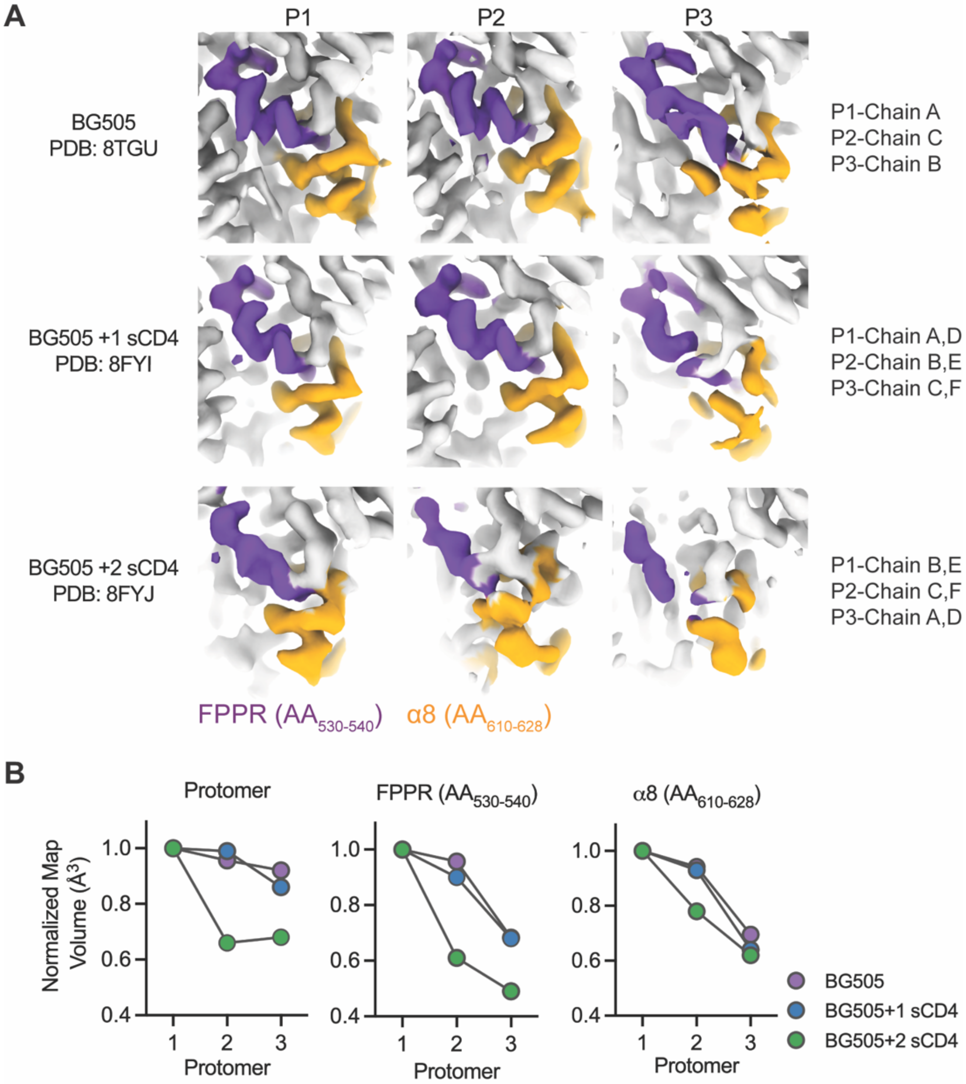


**Supplemental Figure 5. Conformational variability of the gp41 FPPR and alpha-8 clasp regions in the BG505 Env ectodomain. (A)** Cryo-EM density for the fusion peptide-proximal region (FPPR; residues 530-540, purple) and the alpha-8 helix (residues 610-628, yelllow) shown ligand-free BG505 Env (PDB: 8TGU), single-CD4-bound prefusion closed state (BG505 +1 sCD4, PDB: 8FYI), and two CD4-bound open state (BG505 +2 sCD4, PDB: 8FYJ). Each row shows the three Env protomers (P1-P3) separately, with the corresponding chain assignments indicated. The surrounding Env density is colored gray. Images of cryo-EM maps of ligand-free BG505 Env and 2 sCD4-bound BG505 Env from Figure 3 are shown here for comparison. **(B)** Normalized map volumes for the full protomer, FPPR, and alpha-8 regions across the three protomers in each cryo-EM structure. Values were normalized to protomer 1 within each structural state. Ligand-free BG505 is shown in purple, BG505 +1 sCD4 in blue, and BG505 +2 sCD4 in green.

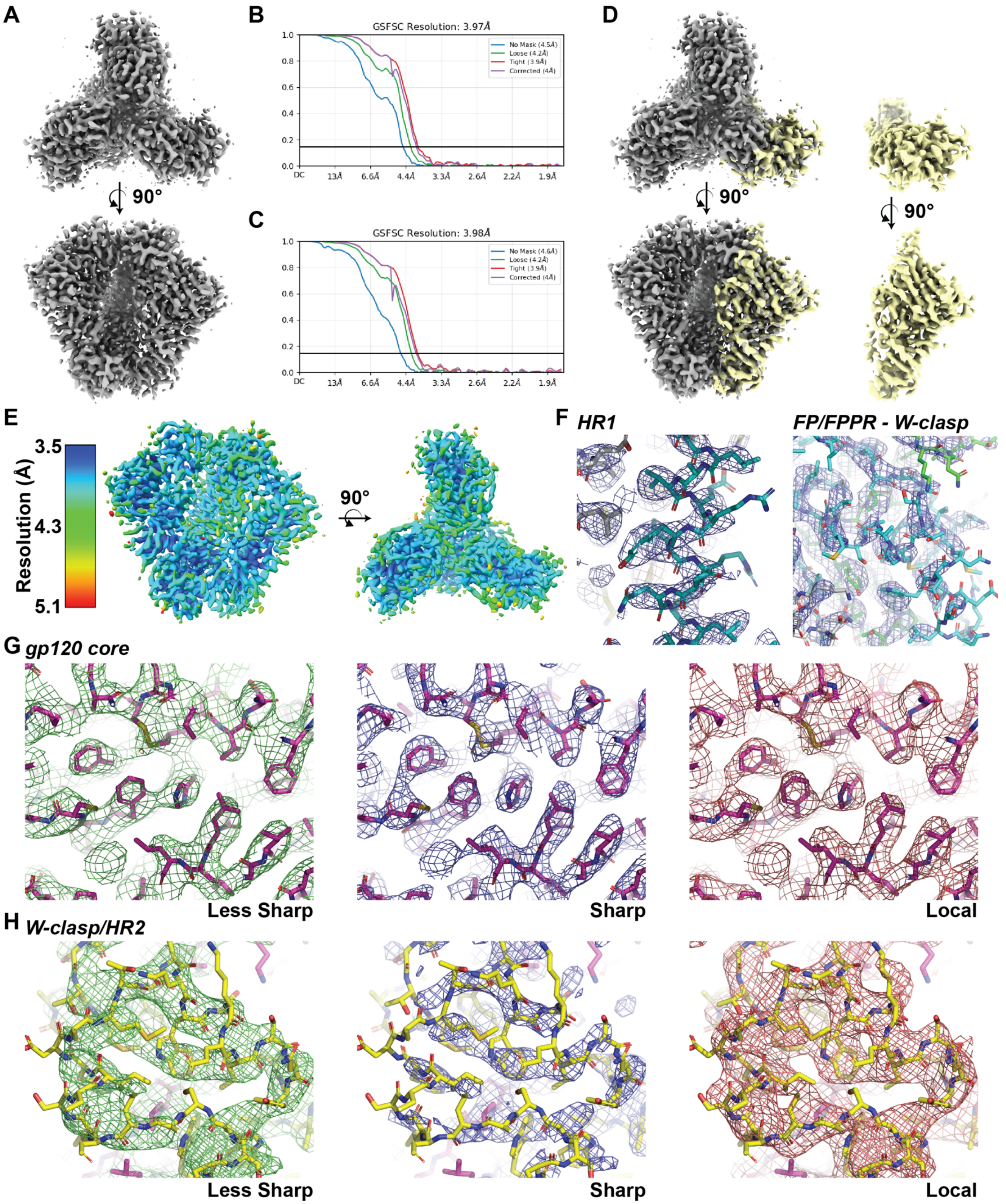


**Supplemental Figure 6. CH505 parent closed-state structure cryo-EM reconstruction. (A)** Sharpened closed-state map viewed from the side and trimer apex. **(B)** FSC plot for the closed-state map. **(C)** FSC plot for single, wide-angle gp41 protomer local-refinement closed-state map. **(D)** Sharpened closed-state map aligned to the local-refinement map viewed from the side and trimer apex. **(E)** Sharpened map colored according to local resolution. **(F and H)** Representative map-to-model fits.

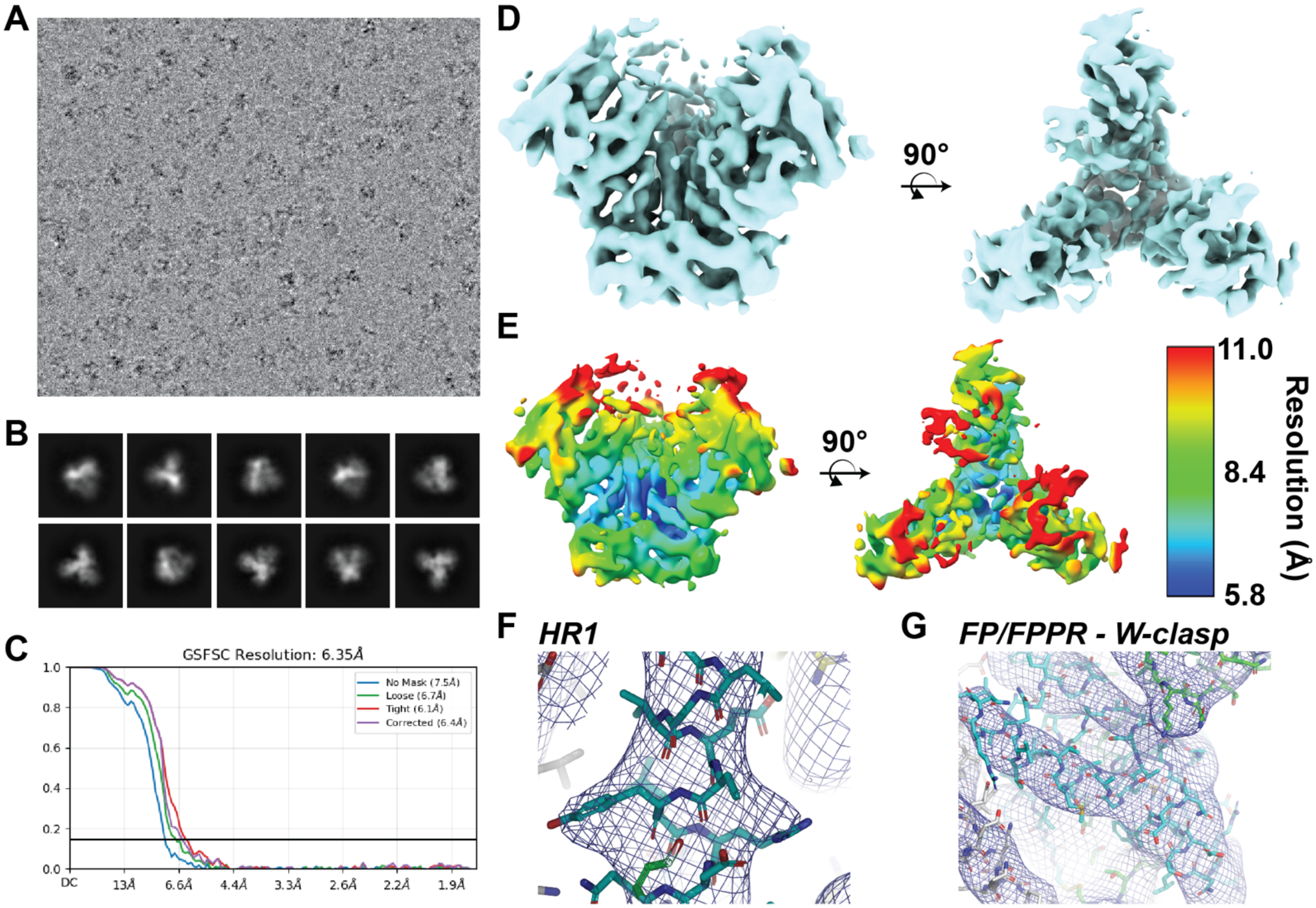


**Supplemental Figure 7. CH505 parent open-state structure cryo-EM reconstruction. (A)** Representative cryo-EM micrograph. **(B)** 2D-class averages for the CH505 parent construct trimer. **(C)** FSC plot for the open-state map. **(D)** Sharpened open-state map viewed from the side and trimer apex. **(E)** Sharpened map colored according to local resolution. **(F and G)** Representative map-to-model fits.

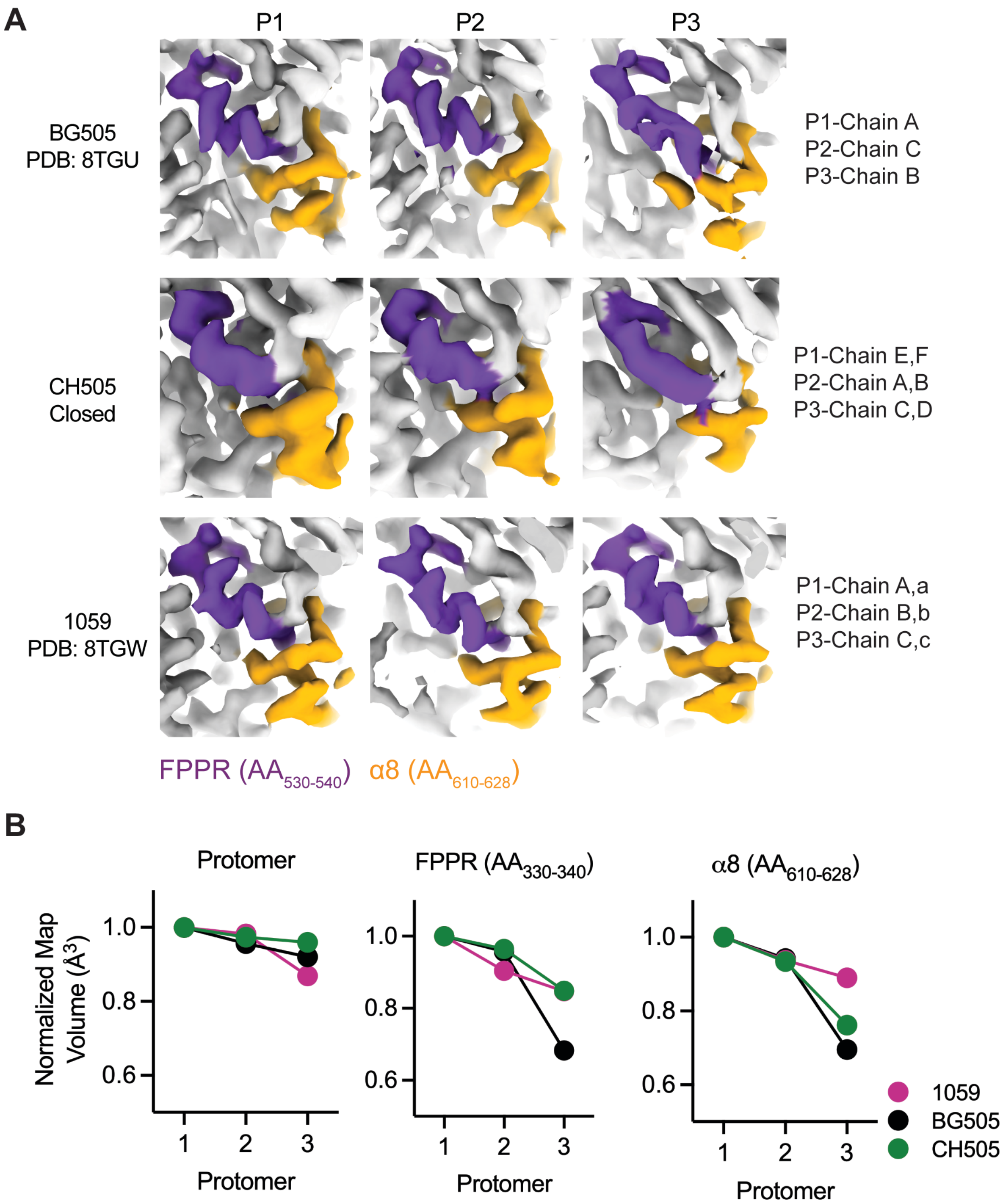


**Supplemental Figure 8. Conformational variability of the FPPR and alpha-8 regions in prefusion-closed Env ectodomain of different HIV-1 isolates. (A)** Cryo-EM density for the fusion peptide-proximal region (FPPR; residues 530–540, purple) and alpha-8 helix (residues 610–628, yellow) in BG505 Env (PDB: 8TGU), 1059 Env (PDB: 8TGW), and closed CH505 Env. The three protomers are shown separately, with the corresponding protomer and chain assignments indicated. Images of cryo-EM maps of ligand-free BG505 Env Figure 3 are shown here for comparison. **(B)** Normalized map volumes for the full protomer, FPPR, and alpha-8 regions. Values were normalized to protomer 1 within each structure.

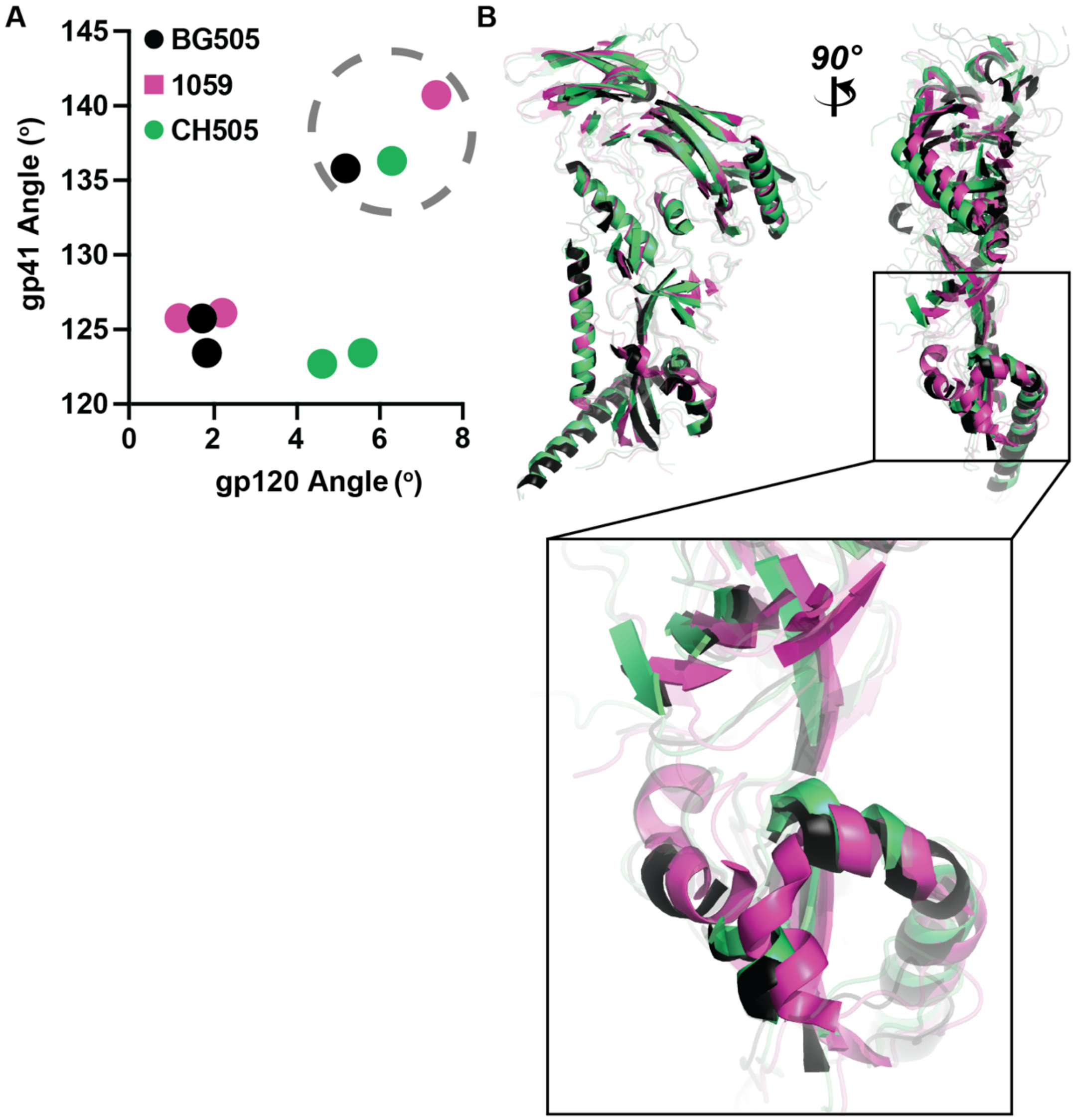


**Supplemental Figure 9. The 1059 gp120 stabilizes the FPPR through direct contact. (A)** gp120 vs. gp41 angles for each protomer. **(B)** Wide-angle gp41 protomer alignment depicting the gp120 displacement relative to the FPPR.

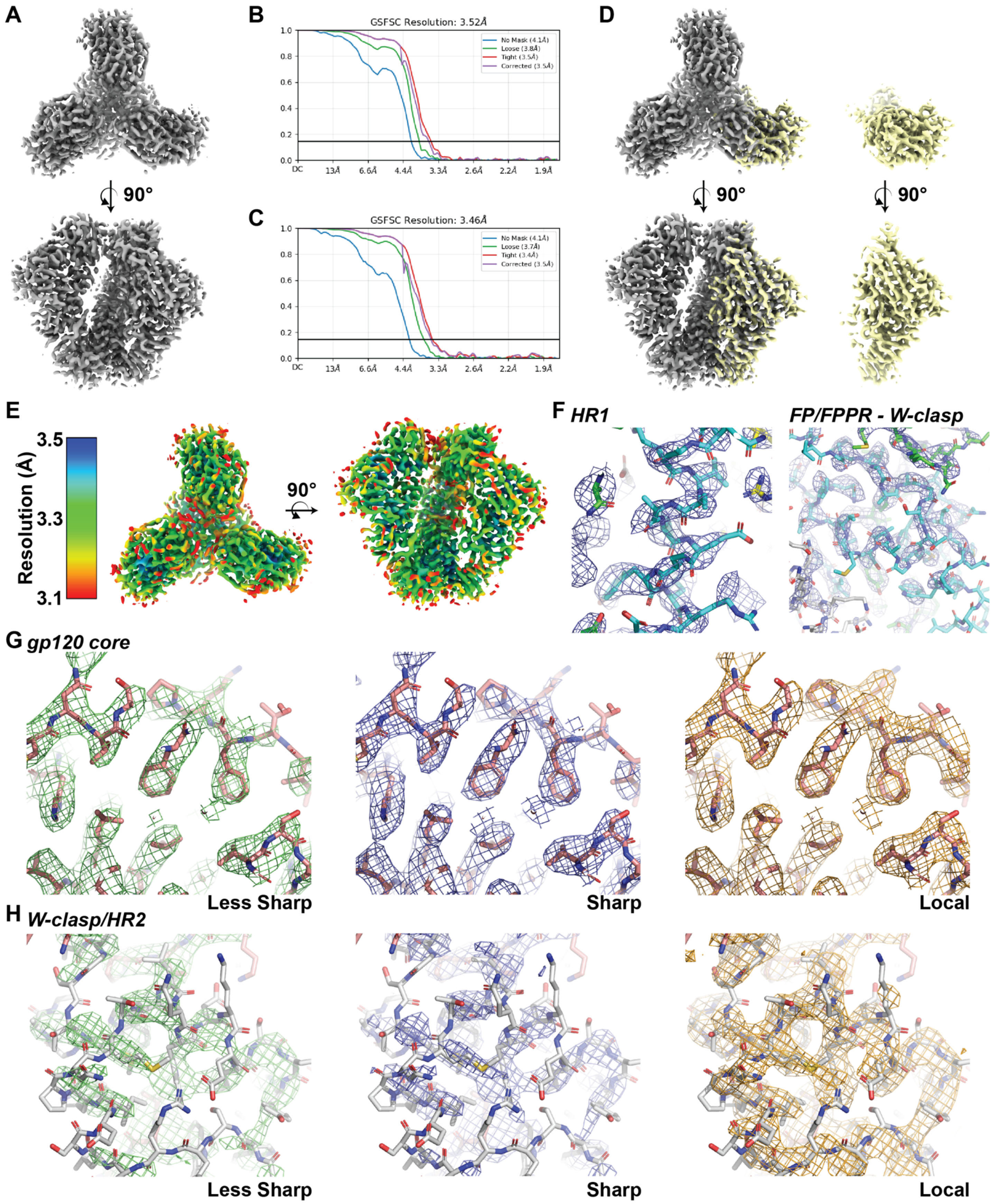


**Supplemental Figure 10. CH505 D624P closed-state structure cryo-EM reconstruction. (A)** Sharpened closed-state map viewed from the side and trimer apex. **(B)** FSC plot for the closed-state map. **(C)** FSC plot for single, wide-angle gp41 protomer local-refinement closed-state map. **(D)** Sharpened closed-state map aligned to the local-refinement map viewed from the side and trimer apex. **(E)** Sharpened map colored according to local resolution. **(F and H)** Representative map-to-model fits.

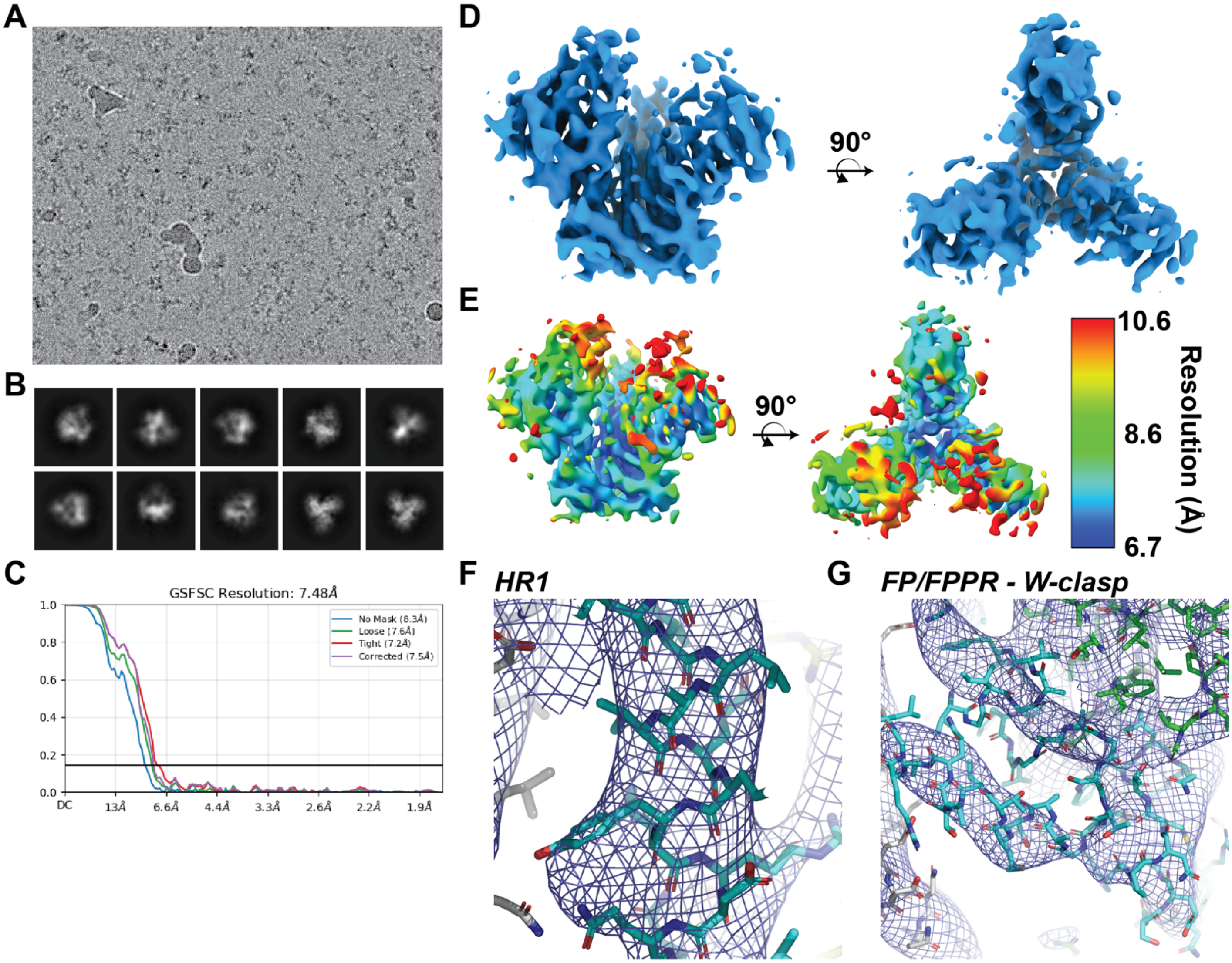


**Supplemental Figure 11. CH505 D624P open-state structure cryo-EM reconstruction. (A)** Representative cryo-EM micrograph. **(B)** 2D-class averages for the CH505 D624P construct trimer. **(C)** FSC plot for the open-state map. **(D)** Sharpened open-state map viewed from the side and trimer apex. **(E)** Sharpened map colored according to local resolution. **(F and G)** Representative map-to-model fits.

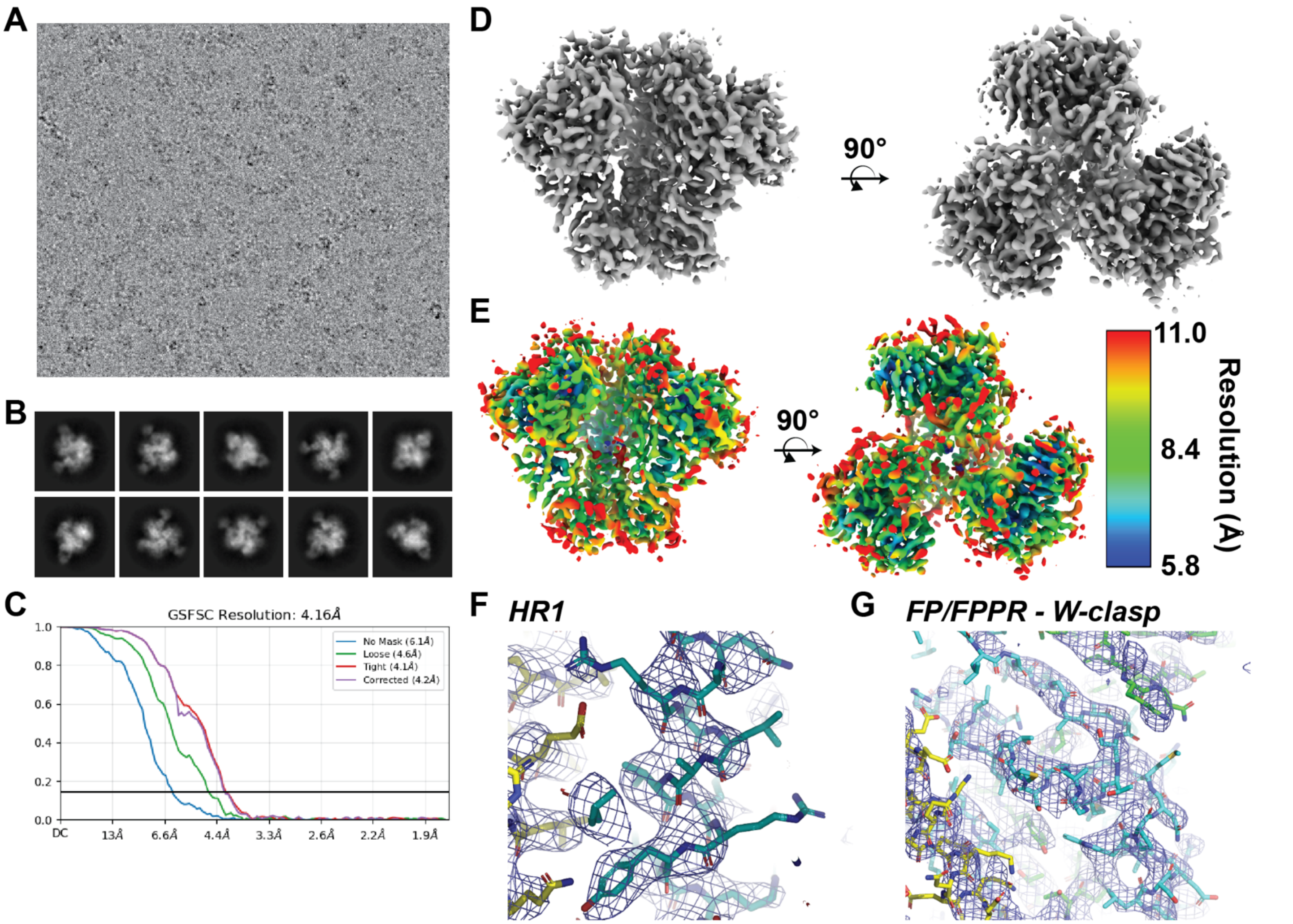


**Supplemental Figure 12. CH235.12 bound CH505 D624P closed-state structure cryo-EM reconstruction.** **(A)** Representative cryo-EM micrograph. **(B)** 2D class averages for the CH235.12-bound CH505 D624P construct trimer. **(C)** FSC plot for the open-state map. **(D)** Sharpened open-state map viewed from the side and trimer apex. **(E)** Sharpened map colored according to local resolution. **(F and G)** Representative map-to-model fits.

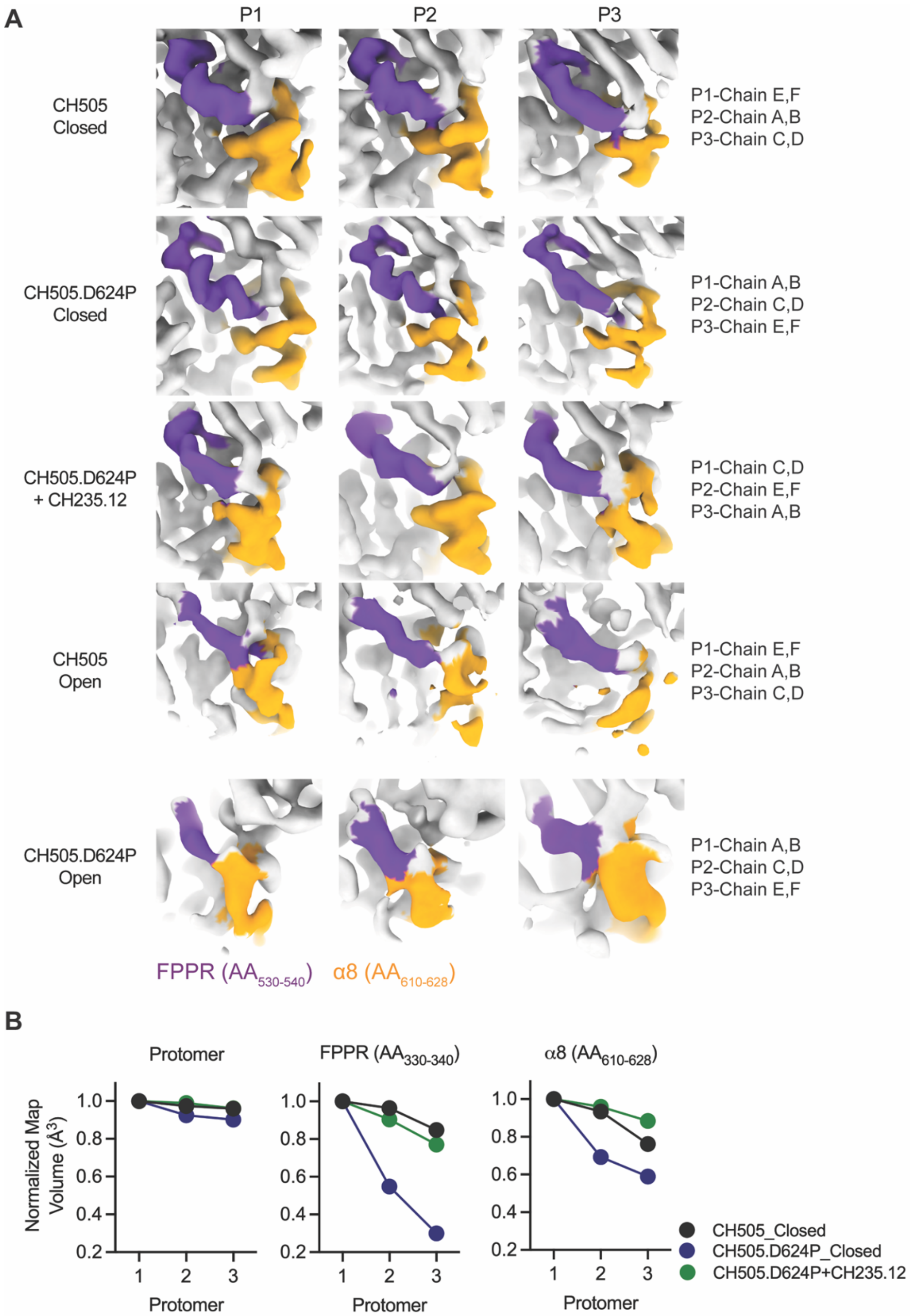


**Supplemental Figure 13: Conformational variability of the FPPR and alpha-8 regions in CH505 Env trimers. A)** Cryo-EM density for the fusion peptide-proximal region (FPPR; residues 530–540, purple) and alpha-8 helix (residues 610–628, yellow) in pre-fusion closed states (CH505, CH505.D624P, and CH235.12-bound CH505.D624P) and pre-fusion open states (CH505 and CH505.D624P). The three protomers are shown separately, with the corresponding protomer and chain assignments indicated. **B)** Normalized map volumes for the full protomer, FPPR, and alpha-8 regions. Values were normalized to protomer 1 within each structure.

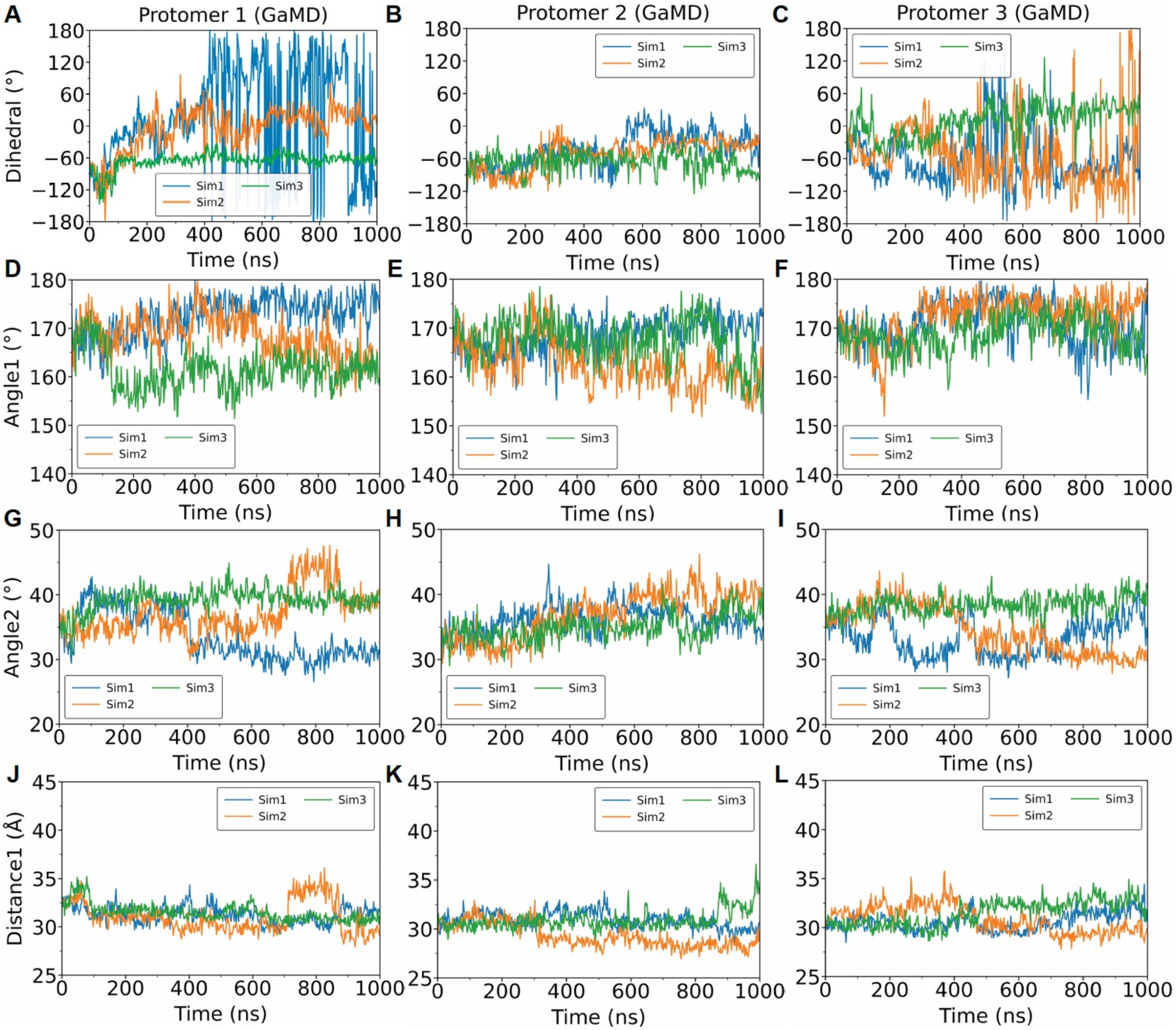


**Supplemental Figure 14. Time series of the dihedral angle, angle 1, angle 2, and distance 1 sampled from the GaMD simulations. (A-C)** Time series of the dihedral $\varphi$ angles formed by $C_\alpha$ atoms of gp41 residue 571, gp41 residue 596, gp120 KK-centroid, and gp120 center-of-mass (COM) in protomer 1 (**A**), 2 (**B**), and 3 (**C**) in the SOSIP. **(D-F)** Time series of the angles formed by $C_\alpha$ atoms of gp120 COM, gp120 KK-centroid, and gp41 residue 596 in protomer 1 (**D**), 2 (**E**), and 3 (**F**) in the SOSIP. **(G-I)** Time series of the angles formed by $C_\alpha$ atoms of gp120 KK-centroid, gp41 residue 596, and gp41 residue 571 in protomer 1 (**G**), 2 (**H**), and 3 (**I**) in the SOSIP. **(J-L)** Time series of the distances between $C_\alpha$ atoms of gp120 COM and gp41 residue 571 in protomer 1 (**J**), 2 (**K**), and 3 (**L**) in the SOSIP.

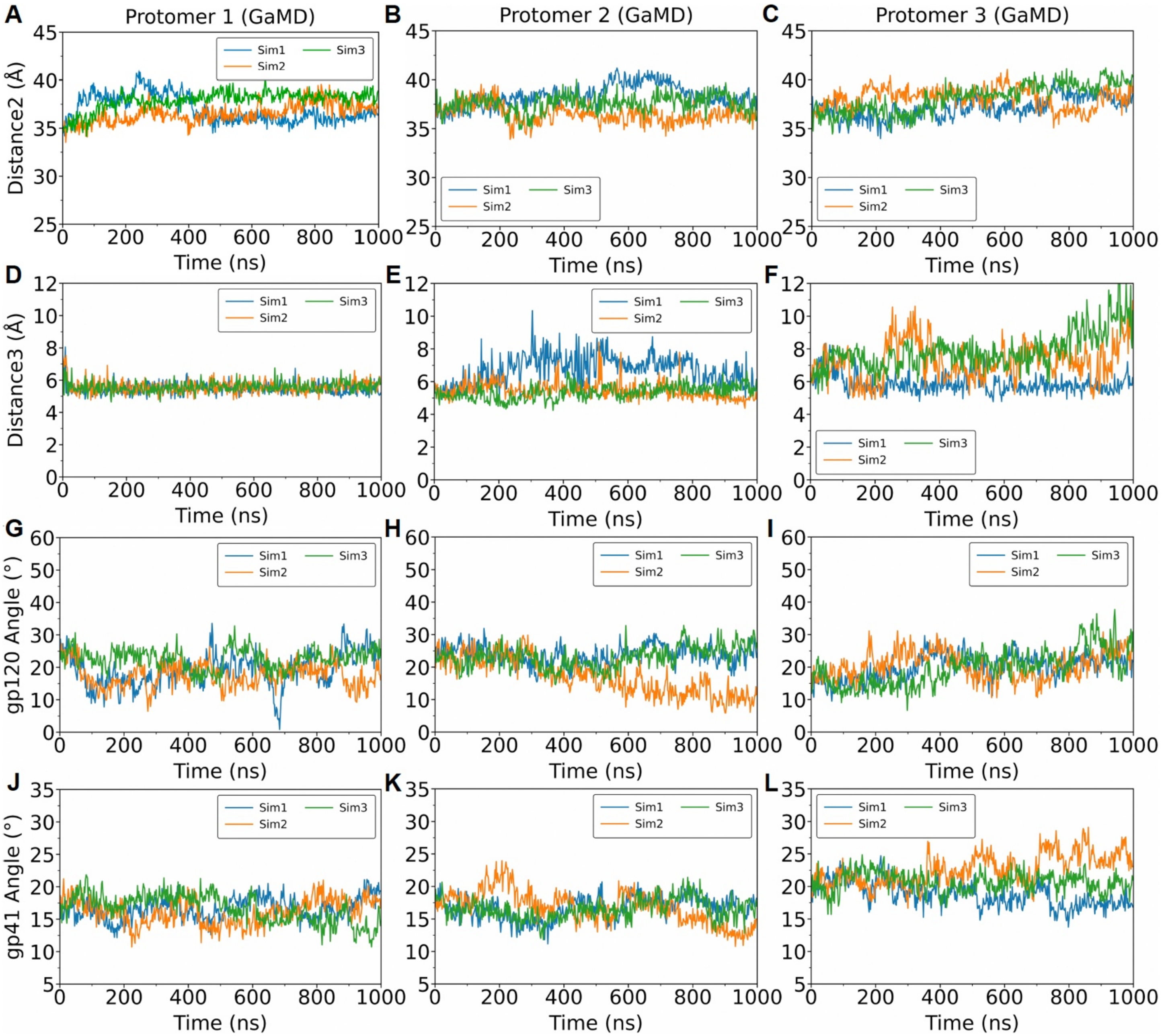


**Supplemental Figure 15. Time series of the distance 2, distance 3, gp120 angle, and gp41 angle sampled from the GaMD simulations. (A-C)** Time series of the distances between $C_{\alpha}$ atoms of gp41 residues 571 and fusion peptide proximal region (FPPR) COM (gp41 residues 528-540) in protomer 1 (**A**), 2 (**B**), and 3 (**C**) in the SOSIP. **(D-F)** Time series of the distances between $C_{\alpha}$ atoms of gp41 residues 530 (of the FPPR) and 623 (of the W-clasp) in protomer 1 (**D**), 2 (**E**), and 3 (**F**) in the SOSIP. **(G-I)** Time series of the tilt angles formed by $C_{\alpha}$ atoms of gp120 COM and the center z-axis drew through the gp41 three-helix bundle in protomer 1 (**G**), 2 (**H**), and 3 (**I**) in the SOSIP. **(J-L)** Time series of the tilt angles formed by $C_{\alpha}$ atoms of gp41 residues 566-596 COM and the center z-axis drew through the gp41 three-helix bundle in protomer 1 (**J**), 2 (**K**), and 3 (**L**) in the SOSIP.

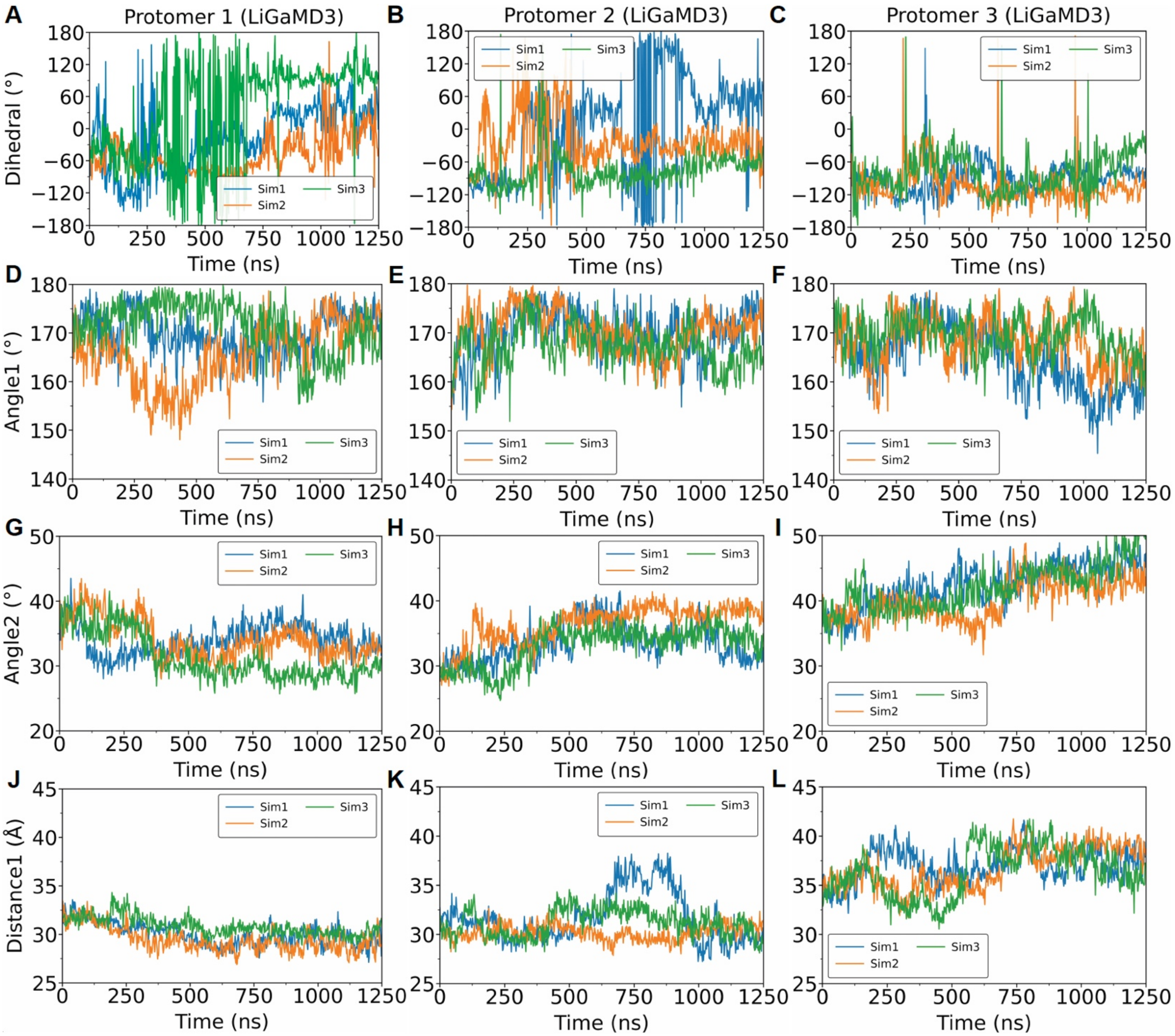


**Supplemental Figure 16. Time series of the dihedral angle, angle 1, angle 2, and distance 1 sampled from the LiGaMD3 simulations. (A-C)** Time series of the dihedral φ angles formed by $C_\alpha$ atoms of gp41 residue 571, gp41 residue 596, gp120 KK-centroid, and gp120 center-of-mass (COM) in protomer 1 (**A**), 2 (**B**), and 3 (**C**) in the SOSIP. **(D-F)** Time series of the angles formed by $C_\alpha$ atoms of gp120 COM, gp120 KK-centroid, and gp41 residue 596 in protomer 1 (**D**), 2 (**E**), and 3 (**F**) in the SOSIP. **(G-I)** Time series of the angles formed by $C_\alpha$ atoms of gp120 KK-centroid, gp41 residue 596, and gp41 residue 571 in protomer 1 (**G**), 2 (**H**), and 3 (**I**) in the SOSIP. **(J-L)** Time series of the distances between $C_\alpha$ atoms of gp120 COM and gp41 residue 571 in protomer 1 (**J**), 2 (**K**), and 3 (**L**) in the SOSIP.

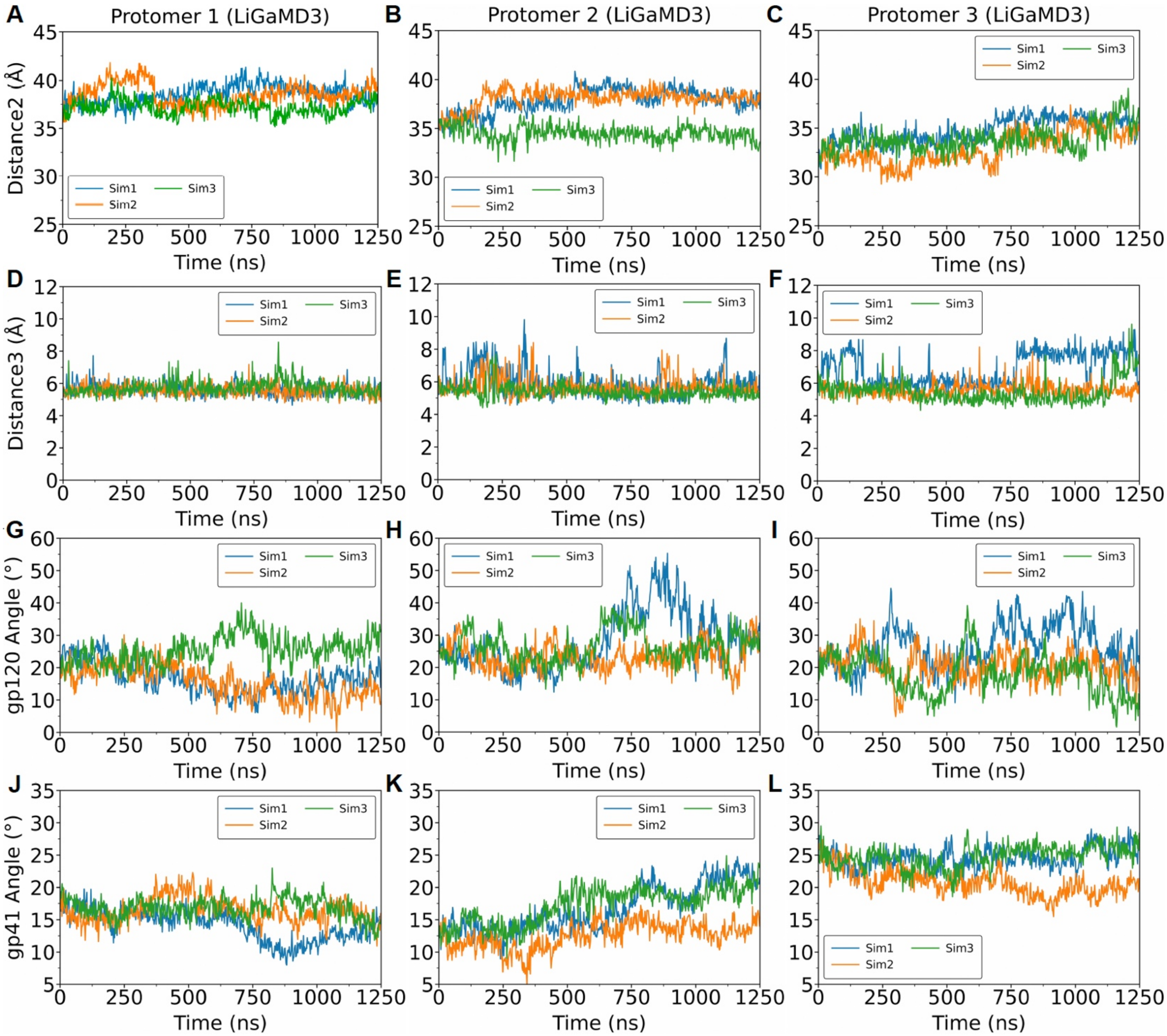


**Supplemental Figure 17. Time series of the distance 2, distance 3, gp120 angle, and gp41 angle sampled from the LiGaMD3 simulations. (A-C)** Time series of the distances between $C_\alpha$ atoms of gp41 residues 571 and fusion peptide proximal region (FPPR) COM (gp41 residues 528-540) in protomer 1 (**A**), 2 (**B**), and 3 (**C**) in the SOSIP. **(D-F)** Time series of the distances between $C_\alpha$ atoms of gp41 residues 530 (of the FPPR) and 623 (of the W-clasp) in protomer 1 (**D**), 2 (**E**), and 3 (**F**) in the SOSIP. **(G-I)** Time series of the tilt angles formed by $C_\alpha$ atoms of gp120 COM and the center z-axis drew through the gp41 three-helix bundle in protomer 1 (**G**), 2 (**H**), and 3 (**I**) in the SOSIP. **(J-L)** Time series of the tilt angles formed by $C_\alpha$ atoms of gp41 residues 566-596 COM and the center z-axis drew through the gp41 three-helix bundle in protomer 1 (**J**), 2 (**K**), and 3 (**L**) in the SOSIP.

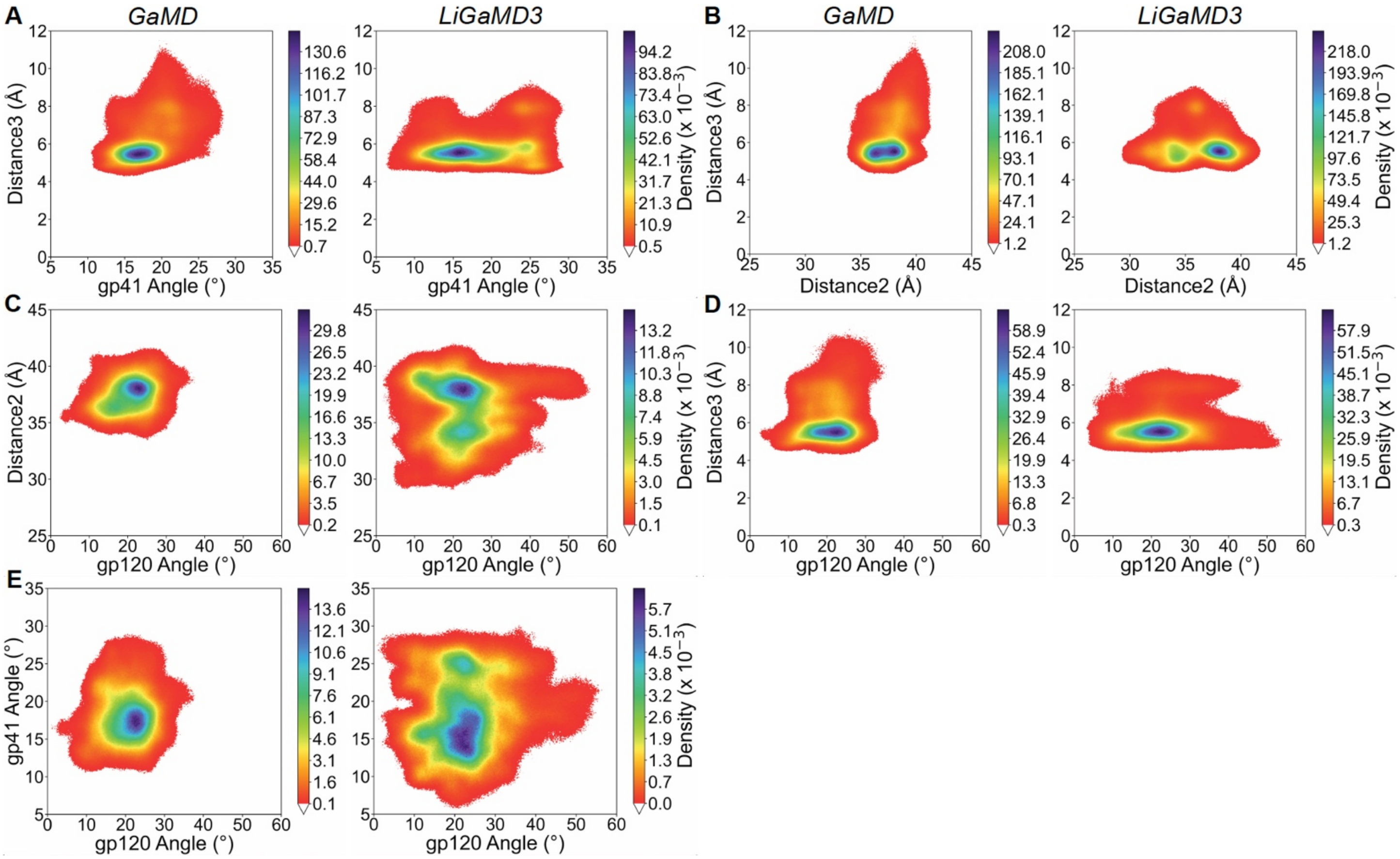


**Supplemental Figure 18. Two-dimensional (2D) Gaussian kernel density estimation (KDE) calculated from GaMD and LiGaMD3 simulations of the SOSIP to monitor the exit of the fusion peptide (FP) from conformational gates. (A)** 2D KDE calculated from the gp41 angle and distance 3. **(B)** 2D KDE calculated from the distances 2 and 3. **(C)** 2D KDE calculated from the gp120 angle and distance 2. **(D)** 2D KDE calculated from the gp120 angle and distance 3. **(E)** 2D KDE calculated from the gp120 and gp41 angles.

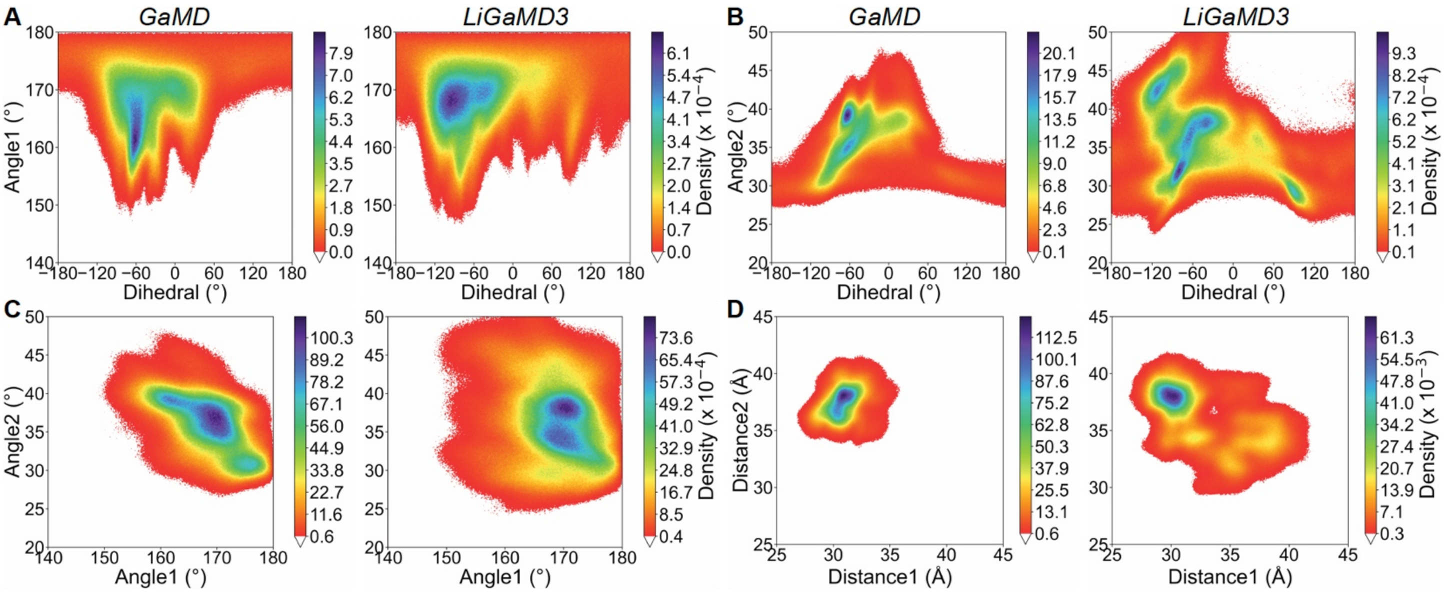


**Supplemental Figure 19. Additional 2D KDE calculated from other collective variables of potential interest in the GaMD and LiGaMD3 simulations of SOSIP. (A)** 2D KDE calculated from the dihedral angle and angle 1. **(B)** 2D KDE calculated from the dihedral angle and angle 2. **(C)** 2D KDE calculated from angles 1 and 2. **(D)** 2D KDE calculated from distances 1 and 2.

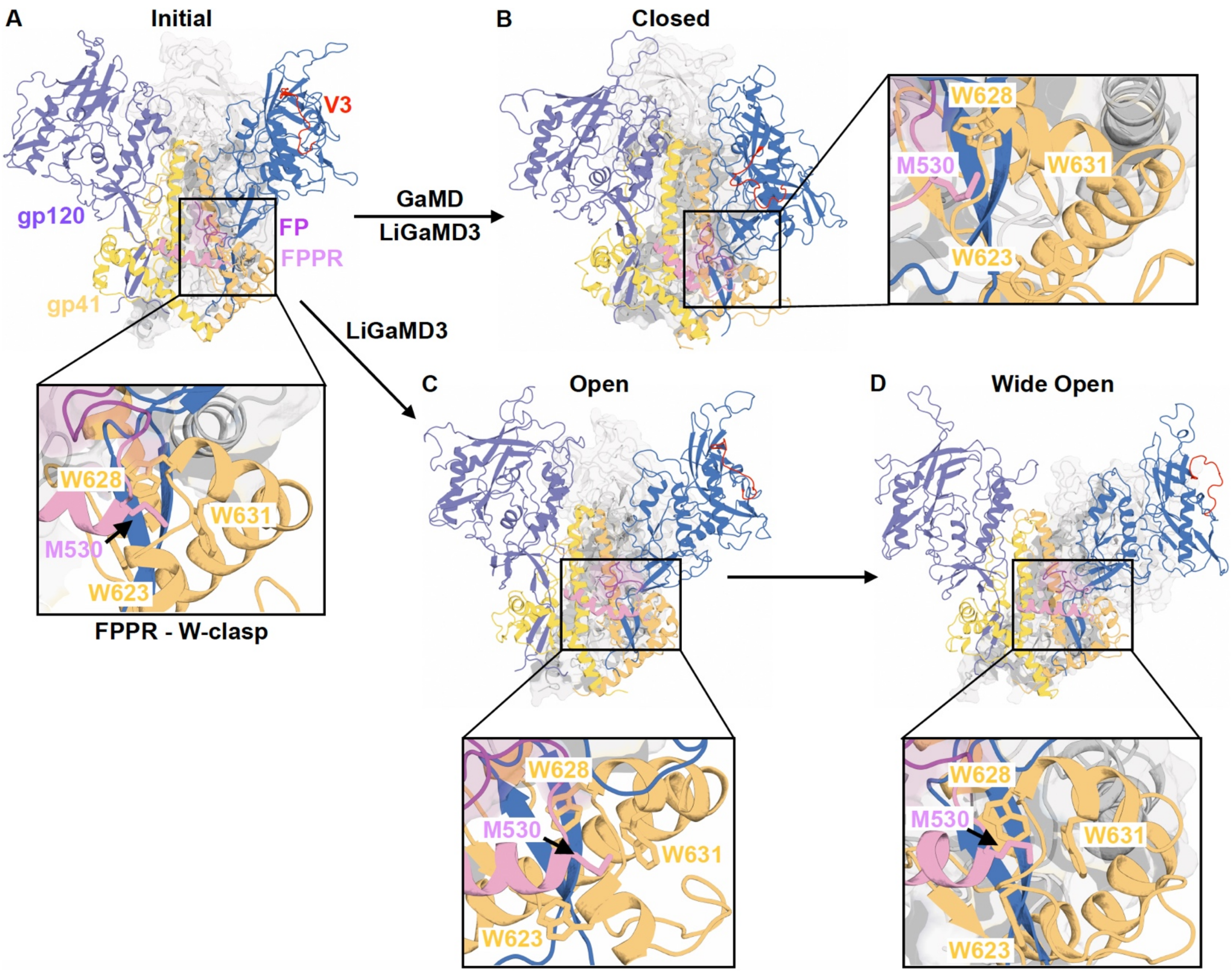


**Supplemental Figure 20. Summary of the conformational transitions of the SOSIP sampled from GaMD and LiGaMD simulations. (A-B)** GaMD simulations only sampled the transition from the initial open (**A**) to closed state of the SOSIP (**B**) due to the absence of CD4 binding, with the FP/FPPR moving downwards and gp120 moving inwards, forcing the FP to become exposed from its original pocket. The closed SOSIP state (**B**) was also sampled during the LiGaMD3 simulations. **(C-D)** LiGaMD3 simulations sampled the transition from the initial open (**A**) to more open (**C**) to widely open (**D**) state of the SOSIP, where the gp120 moved outwards and W-clasp moved outwards, allowing the FP/FPPR to move upwards. The gp120 is colored slate/marine, with the V3 loop colored red, gp41 is colored yellow/bright orange, FP is colored magenta, and FPPR is colored pink. The pocket formed by gp41 residues M530 (of FPPR) and W623, W628, and W631 (of the W-clasp) is zoomed in for clarity.

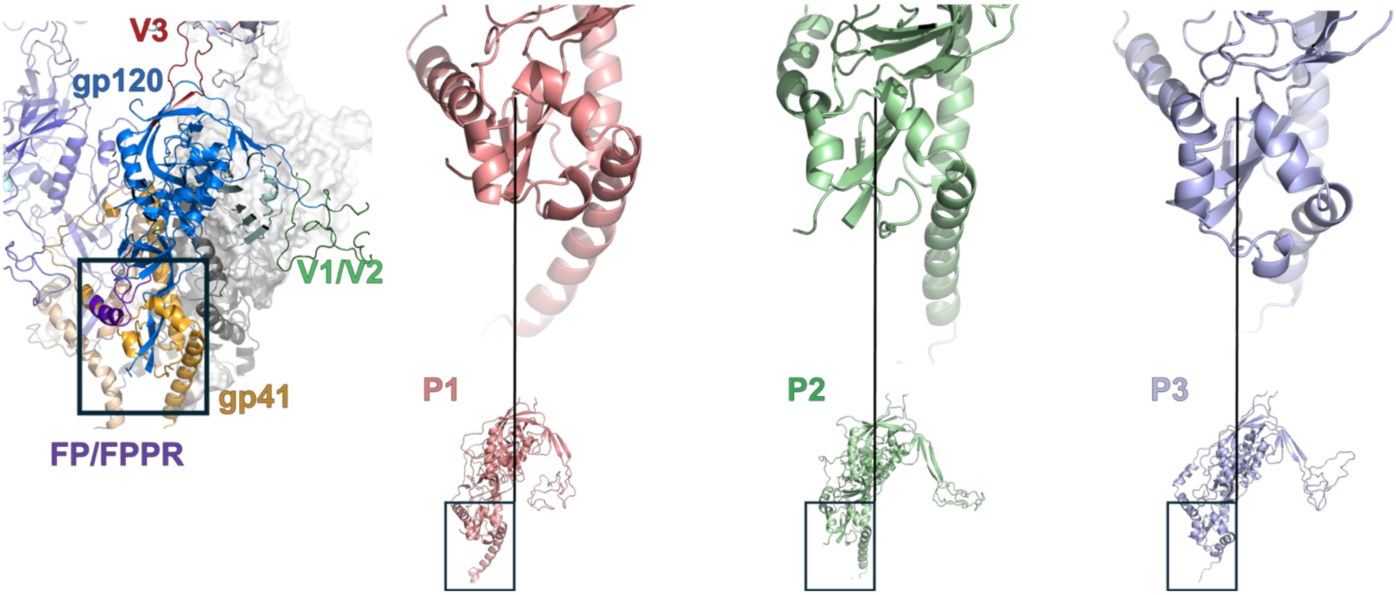


**Supplemental Figure 21. Conformation of the tryptophan-clasp in the full membrane context CD4/CCR5 bound MD simulations.** (left) CH505 Pre-fusion open bound to CD4 and CCR5 highlighting alpha-8/9 and the FP/FPPR region. (right) Zoomed-in views of the helix/sheet motifs in the alpha-8/9 and FPPR regions of the CH505 Pre-fusion open bound to CD4 and CCR5 in Protomer 1 (pink), Protomer 2 (green), Protomer 3 (light purple).

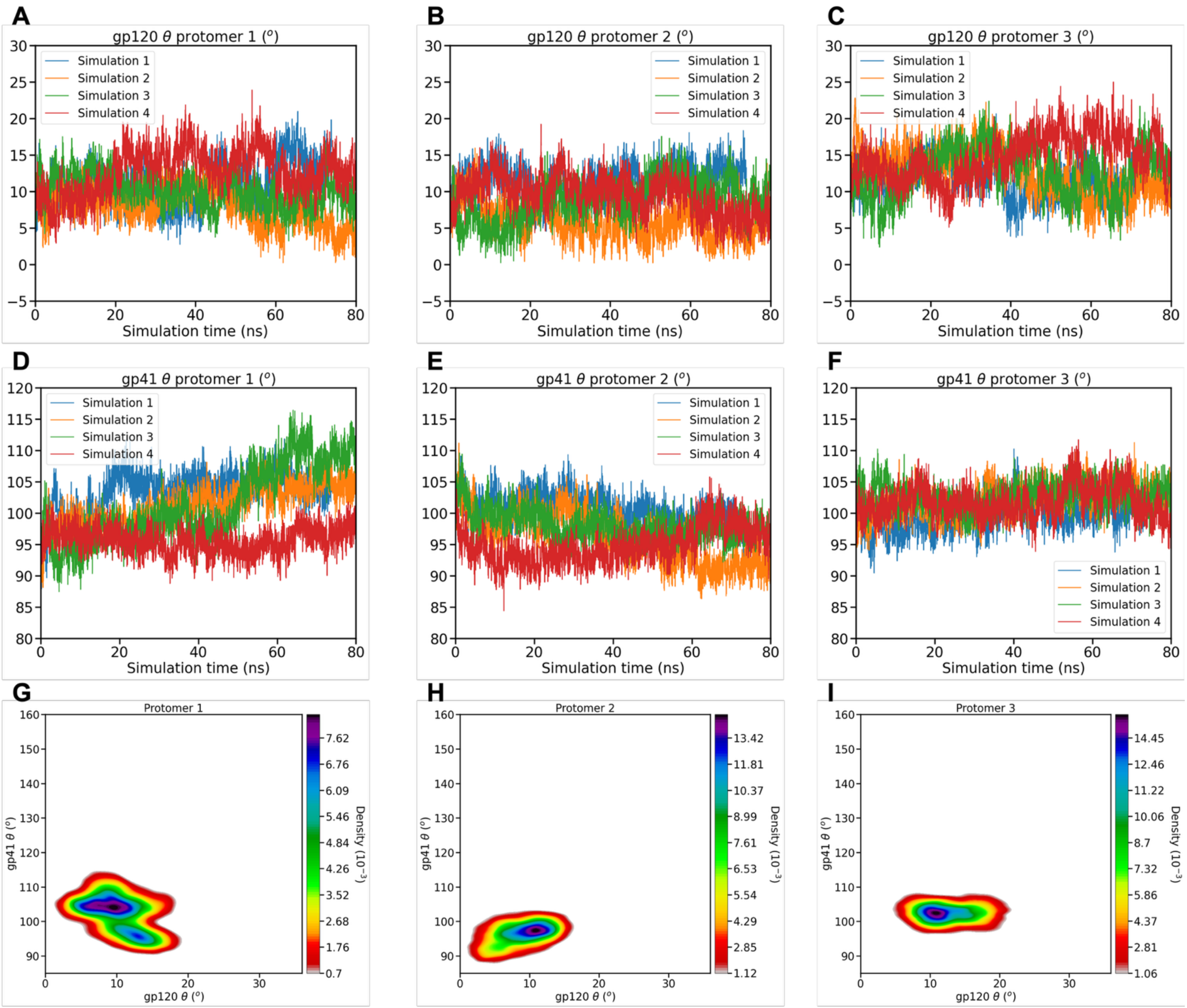


**Supplemental Figure 22. Simulation of fully open Env in complex with CD4 and CCR5 in complex membrane.** Gp120 θ angle of A) protomer 1, B) protomer 2, C) protomer 3 as a function of simulation time. Gp41 θ of D) protomer 1, E) protomer 2, F) protomer 3 as a function of simulation time. Kernel density estimate plot of gp41 θ angle vs gp120 θ angle of G) protomer 1, H) protomer 2, I) protomer 3.

**Supplemental Table 1. HIV-1 gp120 and gp41 sequences used for the MD simulations.**

*gp120 (A32 – V411)*

AKKWVTVYYGVPVWKEATTTLFCASDAKAYDTEVHNVWATHACVPTDPNPQEIVLGNVTENFNMWKNNMV
EQMHEDIISLWDQSLKPCGSSGQACPKVSFEPIPIHYCAPAGFAILKCNSKTFNGSGPCTNVSTVQCTHGIRP
VVSTQLLLNGSLAEEEIVIRSENITDNAKTIIVQLNEAVEINCTRPNNNTRKSIHIGSSGNIRQAHCNISKARWNE
TLGQIVAKLEEQFPNKTIIFNHSSGGDPEIVTHSFNCGGEFFYCNTTPLFNSTWNNTRTDDYPTGGEQNITLQ
CRIKQIINMWQGVGKAMYAPPIRGQIRCSSNITGLLLTRDGGRDQNGTETFRPGGGNMRDNWRSELYKYKV
VKIEPLGIAPTACKRRV

*gp41 (A512 – D664)*

AVGLGAFILGFLGAAGSTMGAASMALTVQARLLLSGIVQQQNNLLRAIEAQQHMLQLTVWGIKQLQARVLAV
ERYLRDQQLLGIWGCSGKIICCTNVPWNDSWSNKTINEIWDNMTWMQWEKEIDNYTQHIYTLLEVSQIQQEK
NEQELLELD

**Supplemental Table 2. Summary of the GaMD and LiGaMD3 simulations performed on the open SOSIP trimer** (obtained from the 5VN3 [1] PDB structure, with CD4 and the 17b antibody removed). Here, $t$ stands for the simulation length, $\sigma_{0P}$, $\sigma_{0D}$, and $\sigma_{0B}$ refer to the upper limits of the standard deviations of the first, second, and third boost potentials, respectively, *iEP*, *iED*, and *iEB* refer to the flags to set the threshold energy for the first, second, and third boost potentials, and $\Delta V$ stands for the added boost potentials.

| System | Method | Replica | $t$ (ns) | $\sigma_{0P}$, *iEP* | $\sigma_{0D}$, *iED* | $\sigma_{0B}$, *iEB* | $\Delta V$ (kcal/mol) |
|---|---|---|---|---|---|---|---|
| Open SOSIP | GaMD | Sim1 | 1000 | 6, 1 | 6, 1 | - | 9.4 ± 3.3 |
| | | Sim2 | 1000 | | | | 9.4 ± 3.4 |
| | | Sim3 | 1000 | | | | 9.7 ± 3.4 |
| | LiGaMD3 | Sim1 | 1250 | 7, 2 | 6, 2 | 6, 1 | 311.3 ± 10.9 |
| | | Sim2 | 1250 | | | | 314.4 ± 11.0 |
| | | Sim3 | 1250 | | | | 310.3 ± 14.2 |

**Supplemental Table 3. Starting values of the collective variables (CVs) monitored for the emergence of the fusion peptides (FPs) for fusogenic competence calculated from the initial structure of the open SOSIP trimer.**

| | **Protomer 1** | **Protomer 2** | **Protomer 3** |
|---|---|---|---|
| **Distance 1 (Å)** | 31.7 | 31.6 | 31.7 |
| **Distance 2 (Å)** | 34.9 | 34.9 | 35.0 |
| **Distance 3 (Å)** | 5.1 | 5.1 | 5.1 |
| **Dihedral (°)** | -100.4 | -94.4 | -97.2 |
| **Angle 1 (°)** | 167.2 | 170.2 | 168.9 |
| **Angle 2 (°)** | 34.0 | 34.0 | 34.0 |
| **gp120 Angle (°)** | 22.1 | 24.7 | 17.9 |
| **gp41 Angle (°)** | 16.4 | 18.1 | 22.4 |

**Supplemental Table 4. Values of the CVs monitored for the emergence of the fusion peptides (FPs) for fusogenic competence calculated from the 3 x 1000 ns GaMD production simulations of the open B41 SOSIP trimer.**

| | **Protomer 1** | **Protomer 2** | **Protomer 3** |
|---|---|---|---|
| **Distance 1 (Å)** | 31.3 ± 0.1 | 30.4 ± 0.5 | 31.0 ± 0.3 |
| **Distance 2 (Å)** | 37.1 ± 0.4 | 37.5 ± 0.6 | 37.8 ± 0.4 |
| **Distance 3 (Å)** | 5.5 ± 0.1 | 5.9 ± 0.4 | 7.0 ± 0.6 |
| **Dihedral (°)** | -20.0 ± 23.3 | -54.2 ± 6.8 | -36.9 ± 22.9 |
| **Angle 1 (°)** | 167.4 ± 3.3 | 166.3 ± 1.8 | 170.6 ± 1.0 |
| **Angle 2 (°)** | 36.6 ± 1.6 | 36.0 ± 0.6 | 35.4 ± 1.5 |
| **gp120 Angle (°)** | 19.9 ± 1.5 | 21.3 ± 2.1 | 20.2 ± 0.3 |
| **gp41 Angle (°)** | 16.5 ± 0.2 | 16.7 ± 0.1 | 20.7 ± 1.1 |

**Supplemental Table 5. Values of the CVs monitored for the emergence of the fusion peptides (FPs) for fusogenic competence calculated from the 3 x 1250 ns LiGaMD production simulations of the open B41 SOSIP trimer.**

| | Protomer 1 | Protomer 2 | Protomer 3 |
|---|---|---|---|
| **Distance 1 (Å)** | 30.1 ± 0.4 | 31.0 ± 0.4 | 36.7 ± 0.2 |
| **Distance 2 (Å)** | 38.1 ± 0.4 | 36.8 ± 1.1 | 33.9 ± 0.5 |
| **Distance 3 (Å)** | 5.6 ± 0.1 | 5.7 ± 0.2 | 6.0 ± 0.4 |
| **Dihedral (°)** | -13.8 ± 22.9 | -30.9 ± 24.6 | -91.2 ± 6.7 |
| **Angle 1 (°)** | 168.9 ± 1.8 | 169.2 ± 1.0 | 167.5 ± 1.4 |
| **Angle 2 (°)** | 33.2 ± 0.9 | 34.4 ± 1.0 | 41.6 ± 0.7 |
| **gp120 Angle (°)** | 19.2 ± 3.1 | 26.0 ± 1.3 | 21.5 ± 2.6 |
| **gp41 Angle (°)** | 15.8 ± 0.8 | 15.4 ± 1.4 | 23.4 ± 1.3 |

**Supplemental Table 6. Changes in residue contacts from the GaMD to LiGaMD3 production simulations of the open B41 SOSIP trimer responsible for the fusion peptide (FP) emergence.** The differences in residue contact frequencies were calculated by subtracting residue contact frequencies calculated from GaMD simulations from those calculated from LiGaMD3 simulations and listed in the parentheses next to the residue contacts. Here, we included residue contacts with absolute differences in contact frequencies ≥ 0.2 (i.e., differences in residue contact frequencies ≤ -0.2 (decreased contacts) or ≥ 0.2 (increased contacts)). A contact definition of ≤ 6Å between $C_{\alpha}$ atoms was used. The HXB2 numbering scheme of gp160 was used to number the residues within the HIV-1 B41 SOSIP.

| B41 SOSIP Protomer | GP120 Residues | GP41 Residues | Residue Contacts |
|---|---|---|---|
| Protomer 1 | K33, K34, V36, G41, V42, P43, Q82, A142, G143, P399, R409, R410, V411 | G514, G516, A517, F518, L520, G521, F522, L523, G524, R579, A582, C605, T606, N607, D612, N620 | G41 – L520 (-0.63), G41 – F522 (-0.48), P43 – G521 (-0.47), V42 – G521 (-0.46), G41 – L523 (-0.43), V42 – L523 (-0.42), P43 – F518 (-0.36), Q82 – G516 (-0.31), P399 – L520 (-0.27), V36 – N607 (-0.22), R410 – T606 (-0.20), R409 – C605 (-0.20) |
| | | | A142 – R579 (0.38), V411 – N607 (0.25), A142 – A517 (0.25), K34 – D612 (0.24), Q82 – G514 (0.23), P43 – G524 (0.23), K33 – N620 (0.23), G143 – A582 (0.23) |
| Protomer 2 | V36, G41, V42, P43, Q82, E83, I84, V85, L86, G87, V89, T90 | L520, G521, L523, G524, A525, A526, G527, Q551, Q552, N553, N554, W610 | G41 – L520 (-0.37), I84 – Q552 (-0.37), E83 – N553 (-0.35), P43 – G521 (-0.35), V42 – G521 (-0.31), V89 – A526 (-0.30), Q82 – N554 (-0.28), E83 – Q552 (-0.28), V85 – Q551 (-0.27), V36 – W610 (-0.21), G41 – L523 (-0.21), T90 – G527 (-0.21), I84 – Q551 (-0.21), V42 – G524 (-0.20) |
| | | | L86 – A526 (0.39), L86 – G527 (0.34), G87 – G527 (0.33), P43 – A526 (0.32), P43 – L523 (0.32), P43 – G524 (0.31), L86 – A525 (0.28) |
| Protomer 3 | V36, G41, V42, V44, W45, P81, Q82, A142, V166, C407, K408, R409, R410 | G516, A517, L520, L523, A578, C605, T606, N607, W610, M629 | A142 – A578 (-0.57), G41 – L523 (-0.41), V36 – W610 (-0.35), V42 – L523 (-0.35), C407 – C605 (-0.33), K408 – C605 (-0.32), R409 – C605 (-0.32), R409 – T606 (-0.30), K408 – N607 (-0.30), K408 – T606 (-0.28), V44 – M629 (-0.22), R409 – N607 (-0.21), G41 – L520 (-0.20) |

| | | | |
|---|---|---|---|
| | | | P81 – G516 (0.30), Q82 – G516 (0.27), Q82 – A517 (0.25), W45 – M629 (0.23), P81 – A517 (0.22), R410 – N607 (0.22), V166 – A517 (0.21) |

**Supplemental Table 7. Average gp120 and gp41 angle from Pre-fusion Fully Open in complex with CD4 and CCR5 Simulations calculated from the last half trajectory of each simulation.**

| | gp120 θ | | | gp41 θ | | |
|---|---|---|---|---|---|---|
| | Average | Standard Deviation | Standard Error | Average | Standard Deviation | Standard Error |
| Protomer 1 | 10.3468627 | 3.61223314 | 0.04116789 | 102.238594 | 5.19685001 | 0.05922745 |
| Protomer 2 | 8.81594375 | 3.36188437 | 0.03831472 | 96.2723397 | 3.19311102 | 0.03639124 |
| Protomer 3 | 12.6810617 | 3.66089257 | 0.04172245 | 102.362977 | 2.52305591 | 0.02875476 |
| All protomers combined | 10.6146227 | 3.88714724 | 0.02557721 | 100.291304 | 4.75411618 | 0.03128182 |
